# Biochemical Impact of p300-Mediated Acetylation of Replication Protein A: Implications for DNA Metabolic Pathway Choice

**DOI:** 10.1101/2024.04.22.590612

**Authors:** Onyekachi Ononye, Sneha Surendran, Olivia K. Howald, Athena Kantartzis-Petrides, Matthew R. Jordan, Diana Ainembabazi, Marc S. Wold, John J. Turchi, Lata Balakrishnan

**Affiliations:** Department of Biology, School of Science, Indiana University Indianapolis, Indianapolis, IN, 46202; Department of Pathology, Harvard Medical School, Boston, MA 02115; Department of Medicine, Indiana University School of Medicine, Indianapolis, IN 46202; Department of Biochemistry and Molecular Biology, Carver College of Medicine, University of Iowa, Iowa City, IA 52242

**Keywords:** Replication Protein A (RPA), p300, lysine acetylation, single-strand DNA binding, G1/S phase, UV-induced damage

## Abstract

Replication Protein A (RPA), a single-stranded DNA (ssDNA) binding protein, is vital for various aspects of genome maintenance such as replication, recombination, repair and cell cycle checkpoint activation. Binding of RPA to ssDNA protects it from degradation by cellular nucleases, prevents secondary structure formation and illegitimate recombination. In our current study, we identified the acetyltransferase p300 to be capable of acetylating the 70kDa subunit of RPA in vitro and within cells. The acetylation status of RPA was increased specifically during the G1/S phase of the cell cycle and also following exposure to UV-induced damage. Furthermore, we were able to specifically identify RPA directly associated with the replication fork during the S phase and UV damage to be acetylated. Based on these observations, we evaluated the impact of lysine acetylation on the biochemical properties of RPA. Investigation of binding properties of RPA revealed that acetylation of RPA increased its binding affinity to ssDNA compared to unmodified RPA. The improvement in binding efficiency was a function of DNA length with the greatest increases observed on shorter length ssDNA oligomers. Furthermore, the mechanism of acetylated RPAs’ increased affinity for ssDNA was shown to be a function of a slower rate of dissociation compared to the unmodified form of the RPA. Our findings demonstrate that p300-dependent, site-specific acetylation enhances RPA’s DNA binding properties, potentially regulating its function during various DNA transactions.

## INTRODUCTION

Replication protein A (RPA) is a highly conserved heterotrimeric protein in eukaryotes, and is involved in various aspects of DNA metabolism such as DNA replication, repair and recombination ^1^. Present at relatively high concentrations within human cells of ∼1 µM ^2^, it functions as the major single strand DNA (ssDNA) binding protein ^3^. During various DNA transactions, high affinity binding of RPA to ssDNA stabilizes the DNA and protects it from degradation from cellular nucleases ^4^. Stabilization of ssDNA structure by RPA also prevents the formation of stable secondary structures that could impede DNA transactions ^5^. Additionally, RPA serves to function as a platform for the assembly of a multitude of replication and repair associated proteins during various biological events. RPA in eukaryotes is composed of 3 subunits: RPA1, RPA2 and RPA3 with molecular weights of 70kDa, 32kDa and 14kDa respectively ^6^. All three subunits are necessary for the formation of a stable and functional RPA complex.

RPA1 has four oligosaccharide/oligonucleotide binding (OB) domains termed DNA binding domains (DBDs) A, B, C and F, while RPA2 and RPA3 have one OB domain each – DBD-D and DBD-E, respectively ^4, 7^. Different RPA DBDs are activated depending on the length of the ssDNA bound to it ^8^. It was previously suggested that RPA shows a sequential mode of interaction with ssDNA; (i) low affinity binding to ∼8 nt ssDNA where DBD-A and DBD-B of RPA1 would interact ^9^, (ii) medium affinity binding to ∼18-20 nt ssDNA with DBD-A, -B and -C of RPA1 interacting, and (iii) high affinity binding to ∼28-30nt ssDNA which implicated all three DBDs of RPA1 (A,B,C) and additionally DBD-D of RPA2 ^9^. However, multiple recent studies have updated the modular binding model to propose a dynamic binding model for RPA to ssDNA, wherein the binding of RPA to substrate is stable, however, it must be dynamically bound such that it easily hands-off the ssDNA substrate to its interacting protein partners ^10^. This model has been further improved in a study using hydrogen-deuterium exchange mass spectrometry (HDX-MS) to show dynamic binding by DBD-A and DBD-B and more stable binding by the TriC core made up of DBD-C, -D and -E ^11^.

During DNA replication, unwinding of the duplex DNA necessitates binding and protection by RPA on both the leading and lagging strand ssDNA templates ^12–14^. On the leading strand, RPA inhibits priming by DNA polymerase alpha/primase (pol α), however, on the lagging strand, the priming function of pol α is stimulated in the presence of RPA ^15^. Additionally, RPA also influences lagging strand synthesis by stimulating the strand displacement activity of DNA polymerase δ (pol δ), which functions to create a 5’ flap structure ^16^. In most circumstances, this structure is recognized, bound and cleaved by flap endonuclease 1 (FEN1) ^17^. However, creation of long 5’ flaps permit stable binding of RPA to displaced flaps preventing FEN1 cleavage ^18, 19^. Processing of RPA-bound 5’ flaps, requires the nuclease/helicase, Dna2, to displace RPA and cleave the flap to a length that is not optimal for RPA rebinding ^20^. FEN1 then cleaves the remainder of the flap, allowing for ligation and maturation of the Okazaki fragments. Thus, RPA functions as a governing switch that dictates the choice of flap processing during Okazaki fragment maturation ^18^.

Furthermore, RPA plays an equally important role in DNA repair as it does in DNA replication. RPA has been implicated in the base excision repair (BER) pathway as it physically interacts with Uracil DNA glycosylase (UNG) ^21–23^, and stimulates the long flap BER pathway ^21^. RPA ssDNA binding activity is also required for nucleotide excision repair (NER) where RPA binds to the undamaged strand of duplex DNA containing bulky DNA adducts ^24–26^. DNA double-strand break (DSB) repair can be accomplished via two main pathways: non-homologous end joining (NHEJ) and homologous recombination (HR) ^27^. RPA is one of the proteins which when phosphorylated, works with phosphorylated Rad51 to find a homologous region for repair of the damaged strands^28^. There is also some evidence implicating other RPA post-translational modifications that could be involved in DNA repair *via* HR, but the exact details are unknown ^29^.

The ssDNA binding function of RPA and its interaction with protein partners are regulated within the cell using a variety of post-translational modifications (PTMs). The most extensively characterized modification is that of RPA phosphorylation, specifically on the N terminus of the RPA2 subunit. Differential phosphorylation of RPA2 occurs during various phases on the cell cycle, with phosphorylation of RPA2 linked to the G1 to S transition ^30^ and dephosphorylation linked to the mitotic phase ^31^. RPA2 is also phosphorylated on exposure to various DNA damaging agents such as hydroxyurea (HU) and UV irradiation suggesting a role for phosphorylation in mitigating the damage response and NER ^6, 32^. Interestingly, lysine residues on RPA1 are also subject to multiple modifications, some which may function as competing changes. For example, RPA1 is known to undergo acetylation, methylation, SUMOylation, ubiquitinylation, and crotonylation. SUMOylation of RPA1 has been demonstrated to aid in Rad51 recruitment to the location of damage to facilitate repair by homologous recombination ^33^. Another PTM observed on all three subunits of RPA is ubiquitination ^34^. Following DNA damage, RPA-ssDNA recruits Ataxia telangiectasia and Rad3-related protein (ATR), ATR interacting protein (ATRIP) kinase and Pre-MRNA Processing Factor 19 (PRP19) complex to trigger phosphorylation and ubiquitination of RPA, which in turn activates ATR-ATRIP and the DNA damage response (DDR) ^34, 35^. As a part of the UV damage response, RPA was shown to undergo lysine acetylation ^36^ which altered the efficiency of NER ^36^. Characterization of RPA1 crotonylation in response to camptothecin (CPT)-induced DNA damage showed that the modification greatly enhanced RPAs interaction with ssDNA ^37^. While lysine mono-methylation of RPA1 has been identified in proteomic analysis, there are no reports characterizing this impact of this modification on biologic or cellular activity.

Our interest in understanding the regulatory mechanism of lysine acetylation on protein function stemmed from the observation that multiple proteins involved in lagging strand maturation are modified by lysine acetylation. Acetylation of FEN1 and Dna2 imparted opposing effects on these proteins with FEN1 showing decreased nuclease activity whereas, Dna2 showed stimulated cleavage function. Since RPA is known to govern the choice of the lagging strand maturation pathway, in our current work we characterized the impact of lysine acetylation on the binding property of RPA. Contrary to previous reports, we found that in addition to PCAF and GCN5, RPA1 is also acetylated by the acetyltransferase p300, both *in situ* and *in vitro* ^36, 38^. The main role of p300 is to act as a transcription coactivator that aids in chromatin remodeling making it accessible for transcription ^39, 40^. Additionally, p300 has also been shown to acetylate many DNA replication and repair proteins including FEN1 ^41^, Dna2 ^41^, WRN ^42^, and DNA polymersase β ^43^. Our work reveals additional lysine acetylation sites on RPA1 than previously reported. We also found an increase in levels of RPA1 acetylation to correspond with the G1/S phase of the cell cycle and observed acetylated RPA directly at the replication fork. Similar to previous reports we observed an increase in RPA acetylation on exposure to UV damage and direct association of acetylated RPA1 with damaged forks. We further analyzed alterations in the dynamic binding of RPA on lysine acetylation and found that acetylation stimulates the binding activity of RPA to bind stably to shorter length ssDNA. The acetylated form of RPA was also slower in dissociating from the ssDNA compared to the unmodified form of RPA in the presence of a high excess of competing substrate. Additionally, the melting and annealing properties of RPA are also influenced by lysine acetylation. Our data suggests that changes to the binding function of acetylated RPA will undoubtedly impact protein-protein interactions, wherein RPA hands-off the substrate to other proteins in either replication or repair pathways.

## RESULTS

### *In vivo and in vitro* acetylation of RPA by acetyltransferase, p300

A study analyzing the acetylation status of proteins in a whole cell extract using high-resolution mass spectrometry (MS) identified lysine (K) acetylation on residues K163, K167 and K259 of hRPA1 ^36^. A subsequent study using serial enrichments of different post-translational modifications found that in addition to RPA1 (K163 and K577), RPA3, the 14 kDa subunit, (K33, K39) was also acetylated ^44^. Other curated proteomic studies have determined K196, K267, K489, K502 to be acetylated on RPA1 and noted on PhosphositePlus website ^45^. In studies using HEK293T cells, the RPA1 subunit was co-transfected with different acetyltransferases, and only GCN5 and PCAF was reported as capable of acetylating RPA1 ^46, 47^. Since numerous proteins in the DNA replication and repair pathway were modified by another acetyltransferase (KAT), p300, and because proteomic analysis identified acetylation signatures on RPA1 that could potentially be linked to p300 related signatures, we tested the ability of p300 to acetylate RPA. To test endogenous cellular RPA acetylation, in the presence and absence of p300, we used the colon cancer p300 wild-type (wt) cell line HCT116 and a p300 deficient cell line (HCT116^p300-^) derived by targeting the exon 2 of the P300 gene. Additionally, we also assessed acetylation of RPA in HCT116^p300-^ cell line, transfected with increasing concentrations (1 µg and 2.5 µg) of plasmid expressing EP300 cDNA (HCT116^p300-^ + EP300). We observed RPA1 to be acetylated in all three cell lines using immunoprecipitation followed by western blot analysis. Acetylation levels of RPA1 was measured by immunoprecipitating proteins using a pan-acetyl lysine, followed by immunoblotting with RPA1 antibody. Comparing the acetylation levels of RPA1, we observed ∼ 2- fold reduction in the acetylation levels of RPA1 in HCT116^p300-^ compared to wtHCT116 cells (Figure 1A). Upon transfection with increasing concentrations of EP300 cDNA into the HCT116^p300-^ cells, we observed a corresponding increase in levels of acetylated RPA1 (Figure 1A). This observation indicates that p300 is fully capable of acetylating endogenous RPA. Though RPA1, showed decreased level of acetylation in the p300 deficient cell line, compared to wild-type, we still observed low levels of RPA1 acetylation suggesting that in addition to p300, other redundant acetyltransferases are capable of acetylating RPA. Expression levels of the other KATs in both the wt and p300 deleted cells are shown in Supplementary Figure 1A. We repeated this experiment in HEK293T cells and observed a p300 associated dose-dependent increase in levels of endogenous RPA1 (Supplementary Figure 1B). We also assessed acetylation status of RPA2 and RPA3 from both HCT116 and HEK293 cells but did not detect any lysine acetylation on these subunits as assessed by western blot analysis (*data not shown*). *In-vitro* acetylation of RPA1 by p300 was further confirmed using autoradiography (Figure 1B) and western blot analysis (Figure 1C). Unmodified RPA and acetylated RPA (modified using the catalytic domain of p300 and ^14^C-acetyl CoA) were subjected to separation by SDS-PAGE gel electrophoresis and stained using Coomassie Brilliant Blue (CBB). We detected all three subunits of RPA on the stained gel (lanes 1, 2, Figure 1B). Autoradiography of the same gel revealed that in addition to the autoacetylation of the catalytic domain of p300, RPA1 and RPA3 (to a small extent) were also acetylated (lane 4, Figure 1B). We also analyzed RPA acetylation by western blot analysis using a pan acetyl lysine antibody and found RPA1 to be robustly acetylated by p300 (lane 3, Figure 1C) and the full-length p300 (lanes 3, 4, Figure 1C) to be autoacetylated. We did not detect acetylation of either RPA2 or RPA3 by western blotting (data not shown).

**Figure 1:**
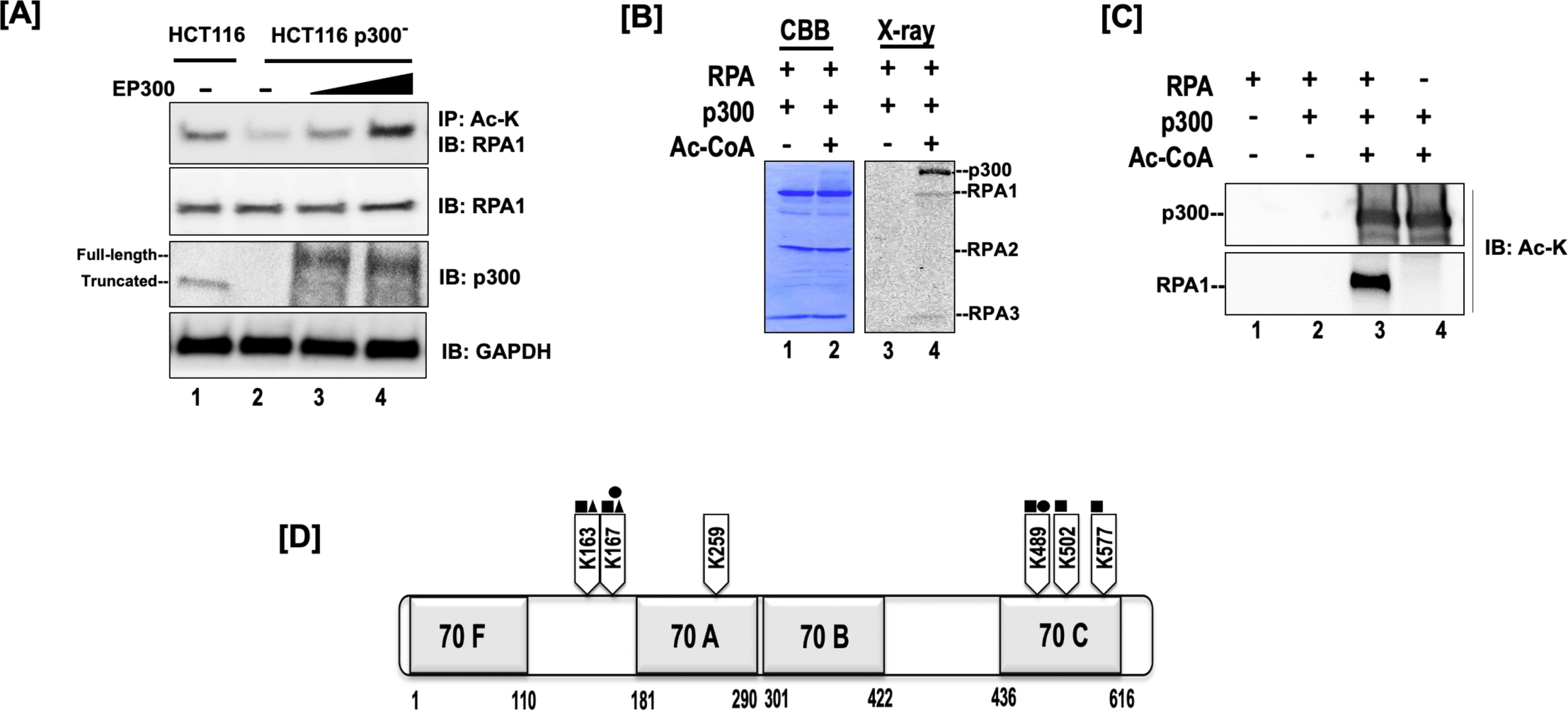
Acetylation of RPA1 Subunit. (A) IP-western blot analysis of RPA1 acetylation in wild-type HCT116, HCT116 p300^-^ and HCT116 p300^-^ cells rescued with EP300 expression. (B) *In vitro* acetylated RPA (Ac-RPA) was subjected to SDS-PAGE analysis and stained using Coomassie brilliant blue (CBB). The same gel was subsequently analyzed by autoradiography (X-ray). (C) In vitro acetylation of RPA and full-length p300 was visualized by Western blot analysis using an anti-acetyl lysine antibody (D) Domains of Replication Protein A Subunit 1. Full length RPA was modified by *in vitro* acetylation using full-length p300 and subject to MS/MS mass spectrometry. Acetylated lysine residues on RPA1 and their positions are denoted. Competing modifications on the same lysine residue previously reported by proteomic studies are indicated, ubiquitination (square), sumoylation (triangle) and methylation (circle).

In order to study the impact of RPA acetylation on its biochemical properties, we used the full-length acetyltransferase, p300, to *in vitro* modify full length RPA containing all three subunits. Sites of acetylation on the p300 modified RPA were determined using tandem mass spectrometry (MS/MS) on tryptic peptides. All peptide masses matched theoretical masses for tryptic peptides for human RPA. Spectra for acetylated peptides showed a mass change of +42 Dalton indicating addition of an acetyl group (Supplementary Figure 1C). Lysine sites that were previously identified to be acetylated on RPA1 in the proteomic analysis of whole cell extracts were also identified in our mass spectrometry analysis (K163, K167, K259, K489, K502 and K577). We were unable to identify any lysine residues on RPA2 or RPA3 that were *in vitro* acetylated by p300. However, this does not imply that these sites are not modified *in vitro*, as the absence of peptides containing these acetylated sites could be due to the poor ionization of the acetylated peptide, or the mass of the peptide being out of range for the set experimental values. Stoichiometric values for the extent of acetylation were not determined in these experiments. Of note, proteomic studies^45^ have also identified lysine residues K163, K167, K489, K502 and K507 on RPA1 to be potential targets for other forms of post-translational modifications such as ubiquitination (U), sumoylation (S) and mono-methylation (M) (Figure 1D), suggesting that within the cell there could be competing or combinatorial PTM on the protein.

### RPA acetylation peaks during the G1/S phase of the cell cycle

Cellular events dictate the PTM status of proteins, altering protein properties (function, stability, localization), and thereby enhance the repertoire of the cellular proteome. Since RPA is vital to DNA replication stability and fidelity, we first determined if cell cycle impacts the acetylation status of RPA. HEK293T cells were synchronized during different cell cycle phases as described in the Methods section and the levels of RPA1 acetylation during the different cell phases was determined. We used a pan-acetyl lysine antibody to immunoprecipitate acetylated proteins from different cell cycle phases, followed by immunoblotting for RPA1 to determine the acetylation status. We also performed this experiment in reverse, wherein we used RPA1 antibody to immunoprecipitate and a pan-acetyl lysine antibody to immunoblot and although we obtained similar results, the blots displayed significant background pixilation (*data not shown*). As shown in Figure 2A, total levels of RPA1 and acetylated RPA1 were altered during different cell cycle phases. Quantitation of AcRPA1 normalized to RPA1 levels, showed that acetylation of RPA1 peaked during the G1/S, followed by S phase of the cell cycle (Figure 2B). Cell synchronizations were confirmed by probing for the differential expression of cyclins during the various phases (Supplementary Figure 2A).

**Figure 2:**
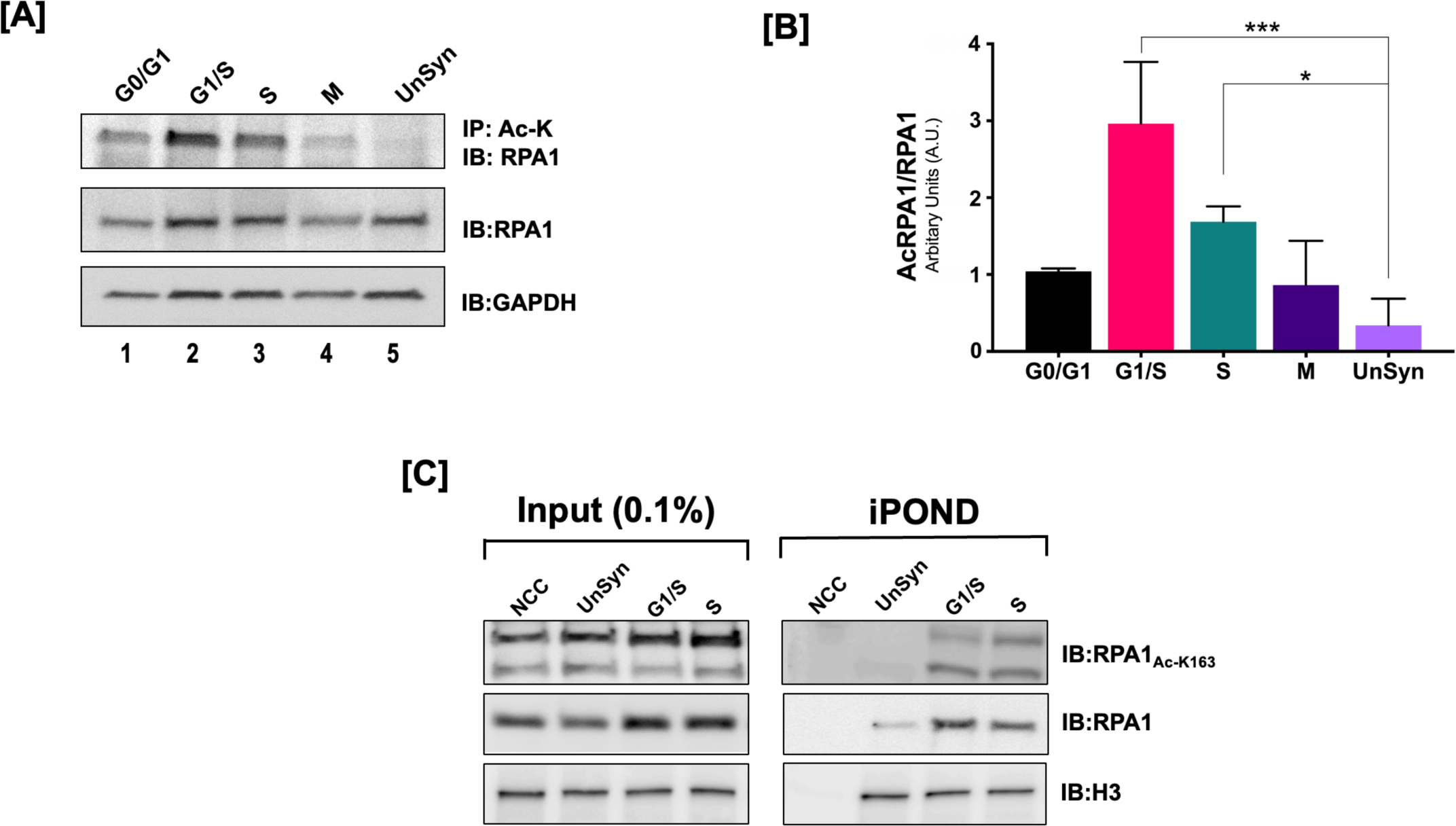
Acetylation of RPA1 is Cell Cycle Dependent: (A) HEK293T cells were synchronized during different phases using either serum starvation or specific chemicals as described in the Materials and Methods. Cell lysates from different cell cycle phases were immunoprecipitated using a pan acetyl-lysine antibody and immunoblotted using RPA1 specific antibody. (B) Graphical representation of change in levels of acetylated RPA1 during different cell cycle phases in HEK293T cells. Statistical significance was assessed by one-way ANOVA * p-value < 0.05, *** p-value < 0.001; (C) HEK293 cells synchronized in various cell phases were subject to an iPOND assay. Lysates from the analysis were evaluated for the presence of acetylated RPA1 on the nascent replication strand. Figures [A] and [C] are representative gels, and the error bars in [B] are for the average of three independent experiments.

While there was a global increase in the acetylation of RPA1 during the G1/S phase, we were particularly interested in determining the acetylation status of RPA1 associated to the replication fork. In order to directly assess acetylation of RPA, we developed an antibody recognizing acetylated K163 of RPA1 (RPA1_Ac-K163_). This antibody eliminated the need to immunoprecipitate RPA followed by immunoblotting using a pan-acetyl antibody. Specificity of the antibody to detect the K163 acetylated residue of RPA1 was confirmed using ELISA (Supplementary Figure 2C). However, while anti-RPA1_Ac-K163_ only detects a 70 kDa band on *in vitro* modified samples, it also detects ∼ a 55 kDa band in cellular extracts, which corresponds to a proteolytic product (Supplementary Figure 2D). This proteolytic product was shown to be stabilized on binding to ssDNA ^48^. Isolation of proteins on naked DNA (iPOND) technique using the RPA1 RPA1_Ac-K163_ revealed a similar trend to our previous result (Figure 2A), wherein, we observed the higher levels of RPA1 acetylation in the G1/S and S phase compared to other phases of the cell cycle (Figure 2C). These results indicate acetylation levels of RPA1 are regulated in response to the cell cycle.

### RPA is acetylated in response to DNA damage repair

Next, we assessed if DNA damage regulates acetylation of RPA, since both Dna2 and FEN1 show increased acetylation on UV damage ^41^. HEK293T cells were exposed to various DNA damaging agents such as hydroxyurea (HU), methane methoxy sulfonate (MMS), ultraviolet radiation (UV), and etoposide (ETP) and the change in acetylation pattern of RPA1 analyzed. DNA damage was confirmed by the presence of markers such as phospho Chk1 (p-Chk1), phospho Chk2 (p-Chk2), phospho p53 (p-p53) and phospho H2AX (p-H2AX) (Supplementary Figure 3). Similar to both Dna2 and FEN1, RPA1 also showed an increase in acetylation on exposure to UV radiation and did not display a detectable increase in acetylation in response to other forms of damaging agents (Figure 3A). RPA1 levels were normalized to one and fold increase in levels of acetylated RPA1 levels calculated and plotted in Figure 3B. Alterations in the phosphorylation of RPA2 serves as a control for known changes in response to DNA damaging agents (Figure 3C). Cells are known to undergo global hyperacetylation in response to either MMS ^49^ or UV ^50^ damage. Similar to the cell synchronization studies (Figure 2D), we were interested in determining if the increase in cellular pools of acetylated RPA1 could be correlated to the RPA that is directly associated with the damaged fork. Using the iPOND assay, we probed for the acetylation status of RPA1 directly associated with the damage fork using anti-RPA1_Ac-K163_ and found that acetylated RPA1 correlated directly with repair of UV damaged forks. Overall, our results suggest that similar to checkpoint kinases that are activated in DNA damage response, RPA1 acetylation could specifically be involved in mediating the UV-induced damage response.

**Figure 3:**
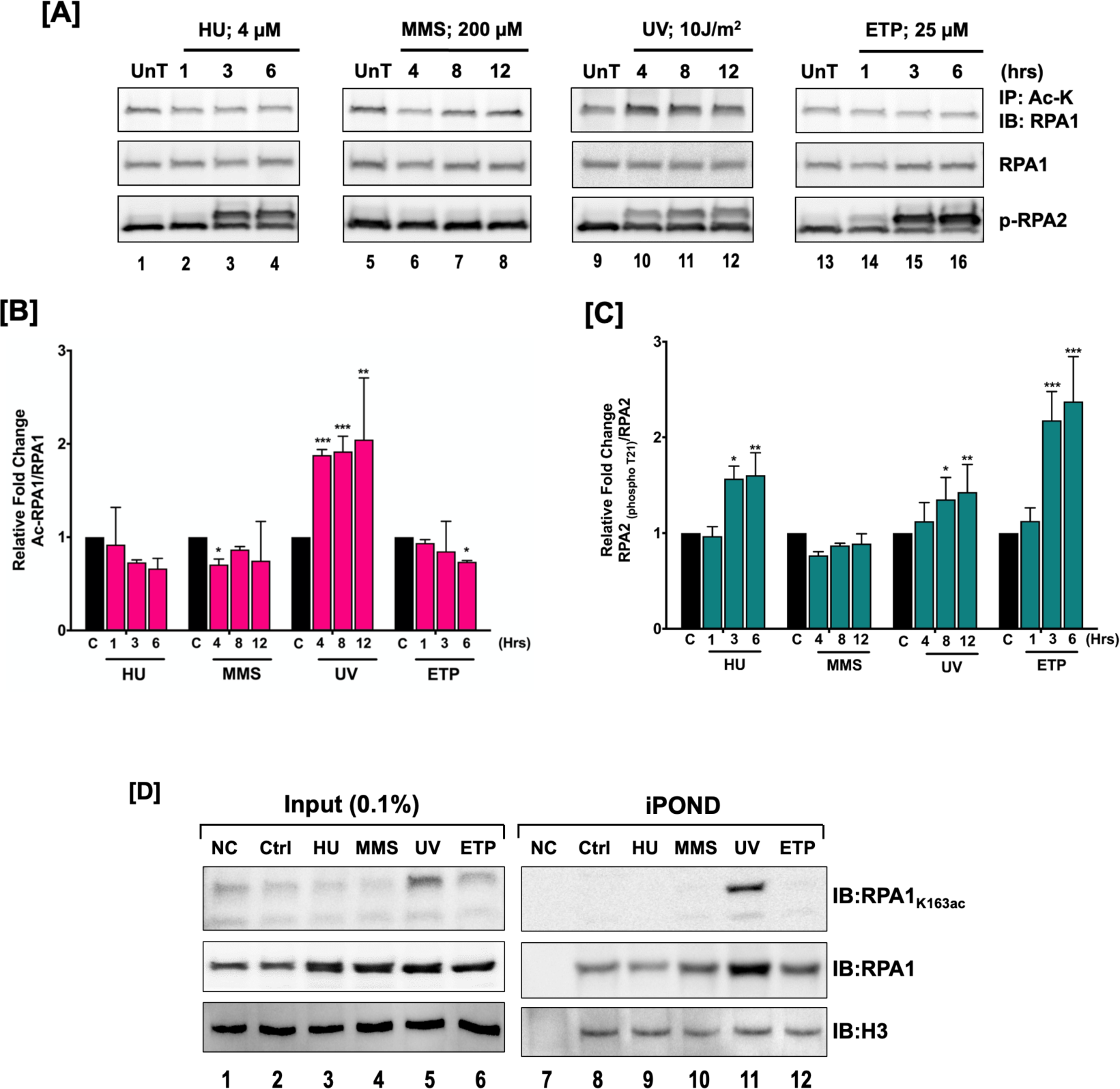
Acetylation is Triggered on UV Damage to the Cells. (A) HEK293T cells were treated with different DNA damaging agents as described in the Materials and Methods. Cell lysates from the different treatments were immunoprecipitated using a pan acetyl-lysine antibody and immunoblotted using RPA1 specific antibody. Graphical representation of (B) change in levels of acetylated RPA1 and of (C) phospho-RPA2 on treatment with different DNA damaging agents in HEK293T cells (D) HEK293 cells treated with different DNA damaging agents (HU and ETP – 6 hours, MMS and UV – 12 hours), were subject to an iPOND assay. Lysates from the analysis were evaluated for the presence of acetylated RPA1 on the nascent replication strand. Figures [A] and [D] are representative gels, and the error bars in [B] and [C] are for the average of three independent experiments. Data is represented as Mean + SEM. Statistical significance was assessed by two-way ANOVA, * p-value < 0.05, ** p-value < 0.005, *** p-value < 0.001.

### Acetylation of RPA increases its ssDNA binding affinity

Binding of RPA to ssDNA is initiated by weak dynamic interactions at lengths of ∼ 8-10 nt involving the A and B domains, while high affinity binding of RPA requires ssDNA to be ≥ 28 nucleotides. We tested the binding efficiencies of unmodified and acetylated RPA on different length oligomers (22, 24, 28 and 32 nt) using electromobility gel shift assays (EMSAs). Based on the length of the ssDNA and known binding properties of RPA, we expected weaker binding to the 22 and 24 nt oligomers compared to high affinity binding to the 28 and 32 nt oligomers. To measure binding, we titrated varying concentrations of RPA (5, 10, 25 nM) in the presence of ssDNA (5 nM) and incubated at 37°C for 10 minutes. Following incubation, the reactions were loaded onto a 6% native gel, electrophoresed, and subsequently analyzed. Binding results indicated that RPA (unmodified and acetylated forms) showed binding to all 4 different length oligomers. Interestingly, irrespective of the length of the oligomer, the acetylated form of RPA (Ac-RPA) showed higher binding efficiency compared to the unmodified form (Um-RPA). However, it is important to note that the fold stimulation in binding of the acetylated form compared to the unmodified form correlated with the lengths of the oligomers. The 22nt oligomer showed the highest fold stimulation in binding by acetylated RPA (compare lanes 6 – 8 to lanes 14-16, 22-24, 30-32, Figure 4A). Smeared pattern of binding to the 22 and 24nt oligomer is consistent with dynamic binding property of RPA to shorter oligonucleotides. Incubation of RPA and p300 in the absence of acetyl coenzyme A showed that it bound similar to the unmodified RPA. The acetyltransferase, p300 was unable to bind to the substrate, suggesting that the observed shift was only due to RPA’s interaction with the substrate (Supplementary Figure 4). The control experiments show that the increased binding efficiency of acetylated RPA was due to lysine modification on the protein and not due to stabilizing interactions with the acetyltransferase.

**Figure 4:**
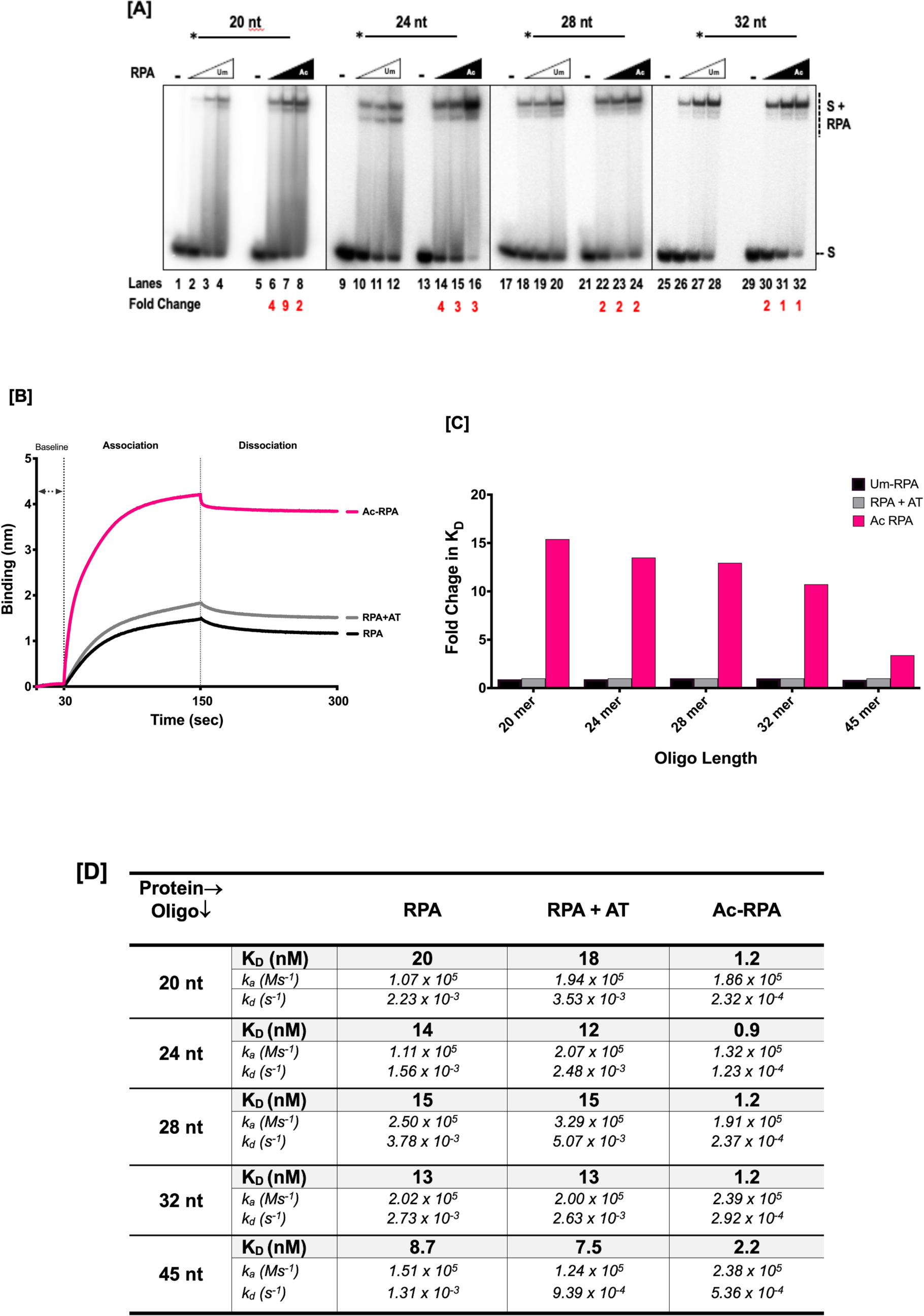
Characterizing the Binding Property of Acetylated RPA. (A) Increase Binding Efficiency of Ac-RPA. Binding efficiency of unmodified and acetylated RPA was studies using EMSA. Five nanomolar of substrate of varying lengths (20, 24, 28 and 32nt) were incubated with increasing concentrations (5, 10, 25 nM) of Um-RPA or Ac-RPA, and the reactions were incubated for 10 min at 37°C and reactions were subsequently separated on a 6% polyacrylamide gel. The labeled substrate is depicted above the gel with the asterisk indicating 5’ of the ^32^P label. The substrate alone and the complexes containing RPA-bound substrate are indicated beside the gel at the right. Fold change in the binding of acetylated RPA compared to the unmodified RPA has been denoted below the lane numbers. (B) Sensorgram obtained using streptavidin biosensor coated with 10 nM 28nt biotinylated oligonucleotide incubated with 160nM RPA, RPA+AT and Ac-RPA. The coating of the biosensor and the association and dissociation curves are shown in the sensorgram. (C) Fold change in binding affinity of RPA+AT and Ac-RPA compared to Um-RPA were calculated. The inverse fold change was then plotted graphically to show the stimulation in binding efficiencies of the Ac-RPA compared to Um-RPA and RPA+AT. (D) Data from the sensorgrams were fit globally to a 1:1 binding model to yield equilibrium dissociation constant (K_D_), association constant (ka) and dissociation constant (kd). Values for K_D_ measurements for different length oligos are listed in the table.

In order to further characterize RPA-ssDNA interactions in real time, we used the label-free biolayer interferometry (BLI) technology to measure protein association and dissociation. Streptavidin biosensors were coated with 10 nM of different length biotinylated oligomers (20, 24, 28, 32 and 45 nt) for a period of 100 seconds and allowed to associate with specific concentrations of either unmodified RPA (RPA), RPA incubated with p300 (RPA+AT) or acetylated RPA (Ac-RPA) for a period of 300 seconds and then moved to a buffer wherein dissociation was measured for a period of 300 seconds. The resulting sensorgram allowed measurement of association and dissociation rate constants (*k_a_* and *k_d_*) and the equilibrium binding constant (K_D_). An example of the sensorgram showing the association and dissociation of 100nM protein (RPA, RPA+AT, Ac-RPA) with a 28 nt oligomer is shown in Figure 4B. Measurements for binding of different concentrations of RPA with different length oligonucleotides were calculated and *k_a_*, *k_d_*, and K_D_ values determined (Figure 4C). Measured binding constants of the unmodified form of RPA agreed with previously reported steady state measurements ^51–53^. For every tested length of oligonucleotide, we found that the acetylated form of RPA had significantly lower K_D_ compared to the unmodified RPA or the RPA + p300 (in the absence of acetyl coenzyme A). While the *k*_on_ of both Um-RPA and Ac-RPA were fairly similar, the off-rate of the Ac-RPA was ∼ 10-fold slower by the Ac-RPA compared to the UM-RPA (Figure 4D). Calculation of the fold change in K_D_ revealed that similar to the EMSA results, fold change in binding constant was the highest for the shortest length oligonucleotide (20 nt) and the lowest for the longest length oligonucleotide (45 nt) measured (Figure 4D). The lower stimulation of binding for the longer length oligonucleotide was expected since RPA is capable of binding to oligonucleotides of this length with high affinity without the need for additional stimulation by acetylation.

### Correlating Sites of Lysine Acetylation on RPA1 to the Increase in Binding Properties

To determine if there was a correlation between increased binding and the number of acetylation sites in the protein’s DBD, we tested mutants of RPA1 subunit containing varying number of DBDs and acetylation sites. The DBD-F mutant contained only DBD-F domain and the linker region with 2 acetylation sites (K163 and K167); the FAB mutant contained DBD-F, DBD-A and DBD-B with 5 acetylation sites (K163, K167, K259, K331 and K379); the A1/A2 mutant contained two DBD-A domains fused together and two acetylation sites (K259,K331) and ΔF-RPA mutant contained all DBDs except DBD-F as well as 7 acetylation sites (K259, K331, K379, K443, K489, K502 and K577) (Figure 5). *In vitro* acetylation of the RPA1 mutants were confirmed both by autoradiography and by tandem mass spectrometry (*data not shown*). Unmodified and acetylated RPA1 mutants were incubated with a 30 nt TAMARA-labeled ssDNA and their binding affinities were analyzed by EMSA. Both the unmodified and acetylated forms of DBD-F mutant did not bind to the substrate. This was an expected result, since the DBD-F does not contribute to ssDNA binding. However, this also confirms that acetylation on sites K163 and K167 do not cause an observable change in the binding property of DBD-F. Additionally, this result further shows that p300 does not complex with DNA to create a gel shift. Acetylation of all of the other mutants (FAB, A1/A2, and ΔF-RPA) showed increased DNA binding compared to their corresponding unmodified forms (Figure 5). Our results suggest that acetylation of one or more lysine residues in RPA1 results in increased ssDNA-binding efficiency, irrespective of the location of the acetylation sites with respect to the DBDs.

**Figure 5:**
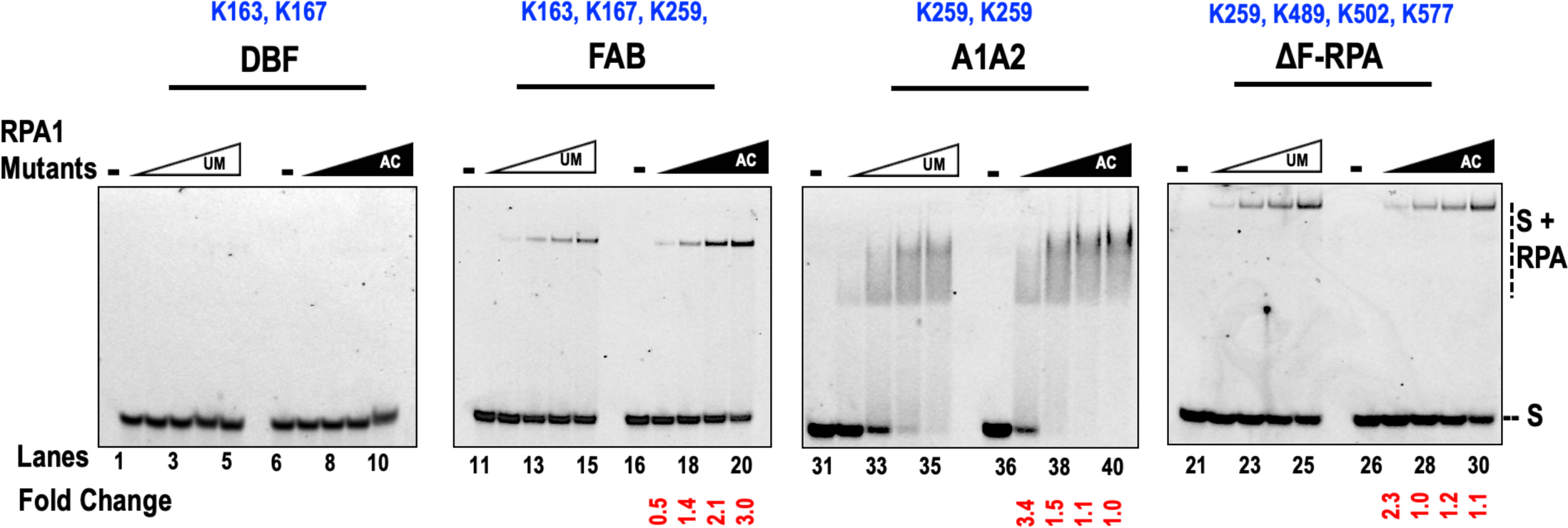
Correlating Number of Acetylated Lysine Sites to Increase in Binding Efficiency. Each lysine site acetylated on the mutant form of RPA1 are indicated above every specific mutant RPA. Binding efficiency of unmodified (Um) and acetylated (Ac) forms of mutant RPA was studied using EMSA. Twenty-five nanomolar of 30 nt 5’TAMARA-labeled ssDNA substrate was incubated in with increasing concentrations (25, 50, 100 and 150 nM) of RPA or Ac-RPA, and the reactions were incubated for 10 min at 37°C and separated on a 6% polyacrylamide gel. The substrate alone and the complexes containing RPA-bound substrate are indicated beside the gel at the right. Fold change in the binding of acetylated RPA compared to the unmodified RPA has been denoted below the lane numbers.

### Acetylated RPA binds more tightly to its substrate compared to the unmodified form

It has been previously shown that ssDNA-bound RPA rapidly exchanges in the presence of free RPA ^10^, however, it can remain stably bound to the ssDNA for many hours ^54^. Given that many biological pathways are dependent on the assembly of RPA on ssDNA and the subsequent hand-off to its interacting protein partners, we were interested in comparing the dissociation of unmodified and acetylated RPA in the presence of a competitor ssDNA substrate. For the competition assays, we chose two ssDNA substrates, a 24 nt and 28 nt substrate. Since only substrates longer than ∼ 28 nt are bound efficiently by RPA, we expected weak binding on a 24 nt ssDNA and tighter binding to 28 nt ssDNA ^55^. We used higher concentration of RPA (unmodified and acetylated) to prebind the 24 nt ssDNA substrate compared to the 28 nt ssDNA substrate, in order to ensure 100% binding of RPA on the shorter length substrate. We pre-bound either Um-RPA or Ac-RPA to a TAMARA-labeled 24 nt or a 28 nt ssDNA and allowed it to incubate for 2 minutes. We then introduced different fold excess (100, 250, 500 and 1000-fold) of a competitor substrate (unlabeled 28 nt substrate) and allowed it to incubate with the reaction for 8 minutes. The reactions were then analyzed using EMSA and the results are graphically represented in Figure 6. The 28 nt competitor unlabeled ssDNA was able to compete off unmodified RPA from both the 24 nt and 28 nt substrate at much lower concentrations compared to the acetylated form of RPA. In the presence of 500-fold excess competitor, nearly 80% of the bound Um-RPA had dissociated from both the 24 nt (black line, Figure 6) and 28 nt substrate (pink line, Figure 6). However, at the same concentration of the competitor, only ∼33% of Ac-RPA was displaced from the 28 nt (pink dotted line, Figure 6) substrate and ∼61% from the 24 nt substrate (black dotted line, Figure 6). Similarly, when all of the bound Um-RPA was displaced in the presence of 1000-fold competitor, 77% of Ac-RPA was displaced from the 24 nt substrate (black dotted line, Figure 6) and 60% from the 28 nt substrate (pink dotted line, Figure 6). This data suggests that the acetylated form of RPA bound more tightly to the substrate and requires a significantly higher amount of competing substrate to be dissociated from its already bound state. This data also correlates with our *k_d_* measurements wherein we observed a slower k_off_ of Ac-RPA compared to Um-RPA (Figure 4D).

**Figure 6:**
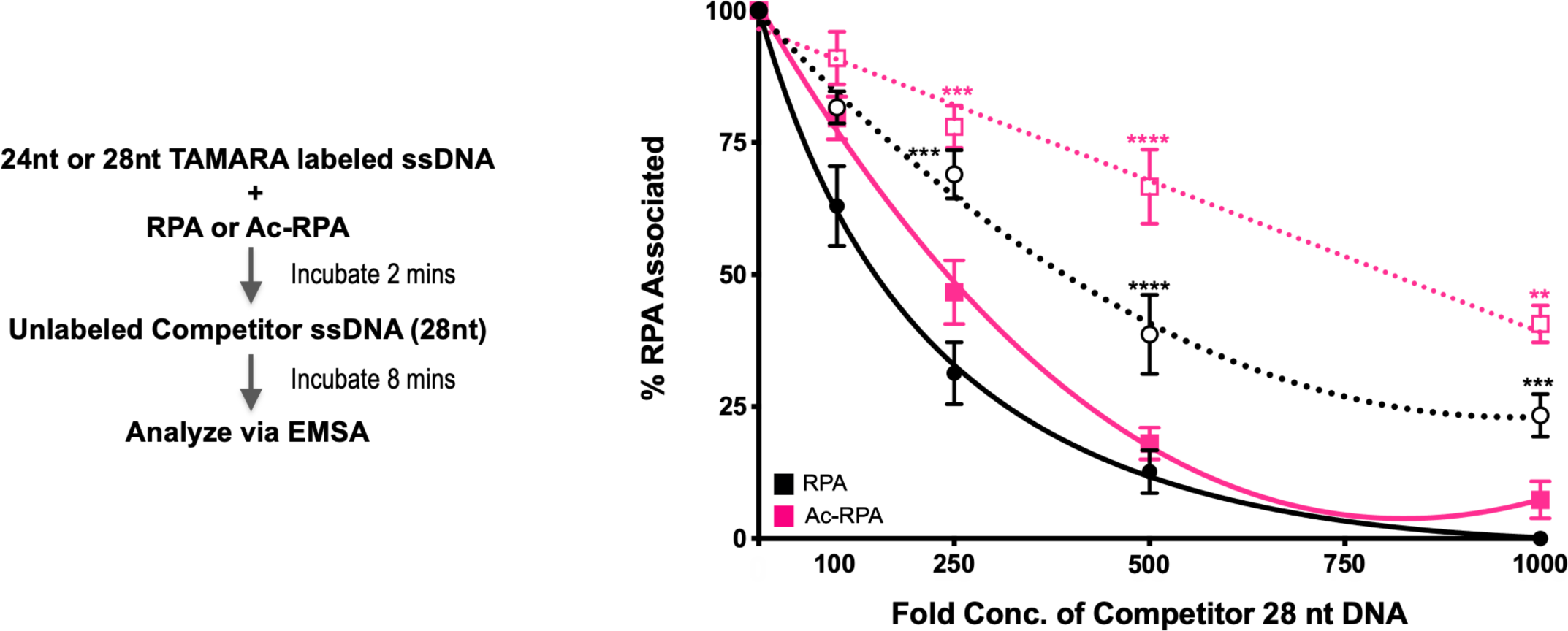
Assaying the Binding Efficiency of Acetylated RPA in Presence of Competing ssDNA. One hundred nanomolar RPA (straight line) or Ac-RPA (dotted line) was pre-incubated with either a 24 nt (black line, round bullet) or 28 nt (pink line, square bullet) TAMARA-labeled ssDNA substrate for a period of 2 minutes at 37°C. To this reaction 100, 250, 500, 1000 - fold excess of cold competitor 28nt ssDNA was added and further incubated for 8 minutes at 37°C and separated on a 6% polyacrylamide gel. Percentage of protein dissociated from the substrate was calculated and graphically plotted to reveal the binding efficiency of RPA and Ac-RPA in the presence of an excess competitor ssDNA. Data is represented as Mean + SEM. Significant two-way ANOVA effects denoted by ** p ≤ 0.05, *** p ≤ 0.001, **** p ≤ 0.0001.

### Assessment of binding inhibition of unmodified and acetylated RPA using small molecule RPA inhibitors

As RPA acetylation is maximal in G1/S phase and enhances DNA binding, we next evaluated the propensity of Ac-RPA to be inhibited by small molecule RPA inhibitors (RPAi) ^56, 57^ that inhibit RPA-DNA binding and exhibit anticancer activity. We utilized two different RPAi, one that is well established (TDRL-551 ^56^) and a second, novel RPAi (DA1-73, manuscript in preparation). We performed RPAi titrations at equimolar Um-RPA and Ac-RPA concentrations where the vehicle controls are ∼17% and ∼77% bound for Um-RPA and Ac-RPA, respectively (Figure 7A and C, lanes 2 and 8). Interestingly, both TDRL-551 (Um-RPA IC50 = 4.46±0.11 µM, Ac-RPA IC50 = 14.12±3.45 µM) and DA1-73 (Um-RPA IC50 = 6.43±0.29 µM, Ac-RPA IC50 = 9.87±2.10 µM) exhibit more potent inhibition of Um-RPA as compared to Ac-RPA (Figure 7B, D). Additionally, as Ac-RPA binds ssDNA more efficiently than Um-RPA (Figure 4), we measured RPA-DNA binding inhibition using Um-RPA and Ac-RPA concentrations that result in 50-60% DNA binding for the vehicle control before titrating RPAi (Supplementary Figure 5A, 5C lanes 2 and 8). Under these conditions, TDRL-551 (Um-RPA IC50 = 4.4±0.9 µM, Ac-RPA IC50 = 4.6±0.7 µM) and DA1-73 (Um-RPA IC50 = 8.4±2.3 µM, Ac-RPA IC50 = 8.6±2.3 µM) inhibit Ac-RPA nearly identically to that of Um-RPA (Supplementary Figure 5B, 5D). Collectively, these data reveal that RPAi can effectively inhibit both Um-RPA and Ac-RPA and may be less effective towards Ac-RPA (Figure 7). However, these differences in inhibition are likely due to the enhanced Ac-RPA binding affinity, as equimolar RPA-DNA complexes are identically inhibited (Supplementary Figure 5).

**Figure 7.**
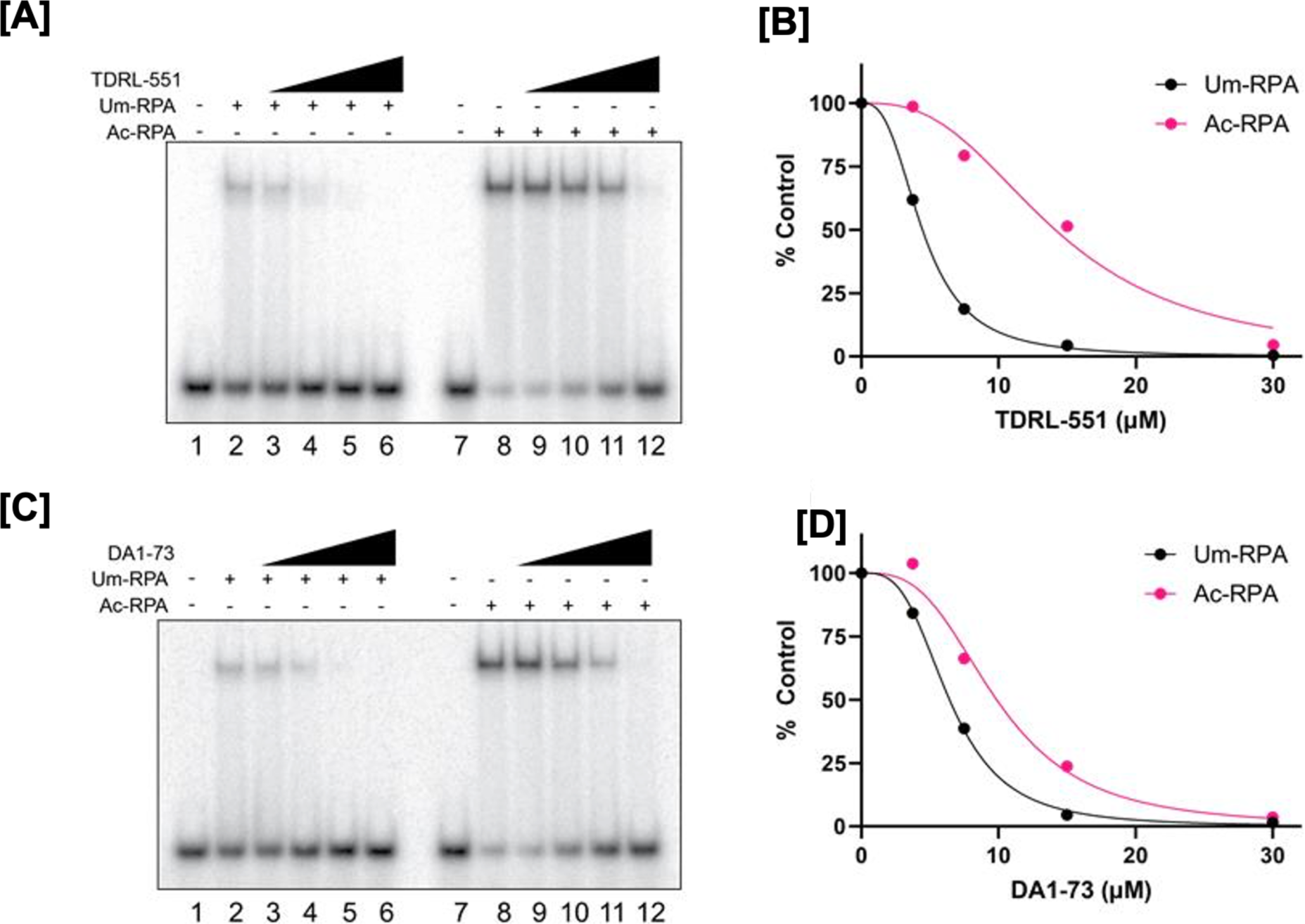
Inhibition of RPA-DNA binding at equimolar RPA-DNA complex. Representative EMSA titration and TDRL-551 (A-B) or DA1-73 (C-D) inhibition of 12.5nM Um-RPA or Ac-RPA. Data in panels B and D are plotted as average ± SEM from three replicates and fit by non-linear regression.

### Strand Melting and Annealing Properties of RPA are Altered by Lysine Acetylation

The inherent binding property of RPA affords it subsidiary functions to either melt or anneal duplex DNA. Since helix destabilization has been linked to RPA1, we were interested in studying the impact of lysine acetylation on this activity of the protein. Alterations in annealing and melting were assessed using the same substrates. In order to characterize the influence of lysine acetylation on RPA annealing, varying concentrations of the protein (10, 25, 50, 75 and 100 nM) were incubated with two complementary ssDNA oligonucleotides at 37°C for 10 minutes. Our results show that there isn’t a significant difference between the percentages of annealed products formed in the presence of both forms of the protein (Figure 8A). However, at a certain point (75nM), Ac-RPA seems to favor the annealing reaction.

**Figure 8:**
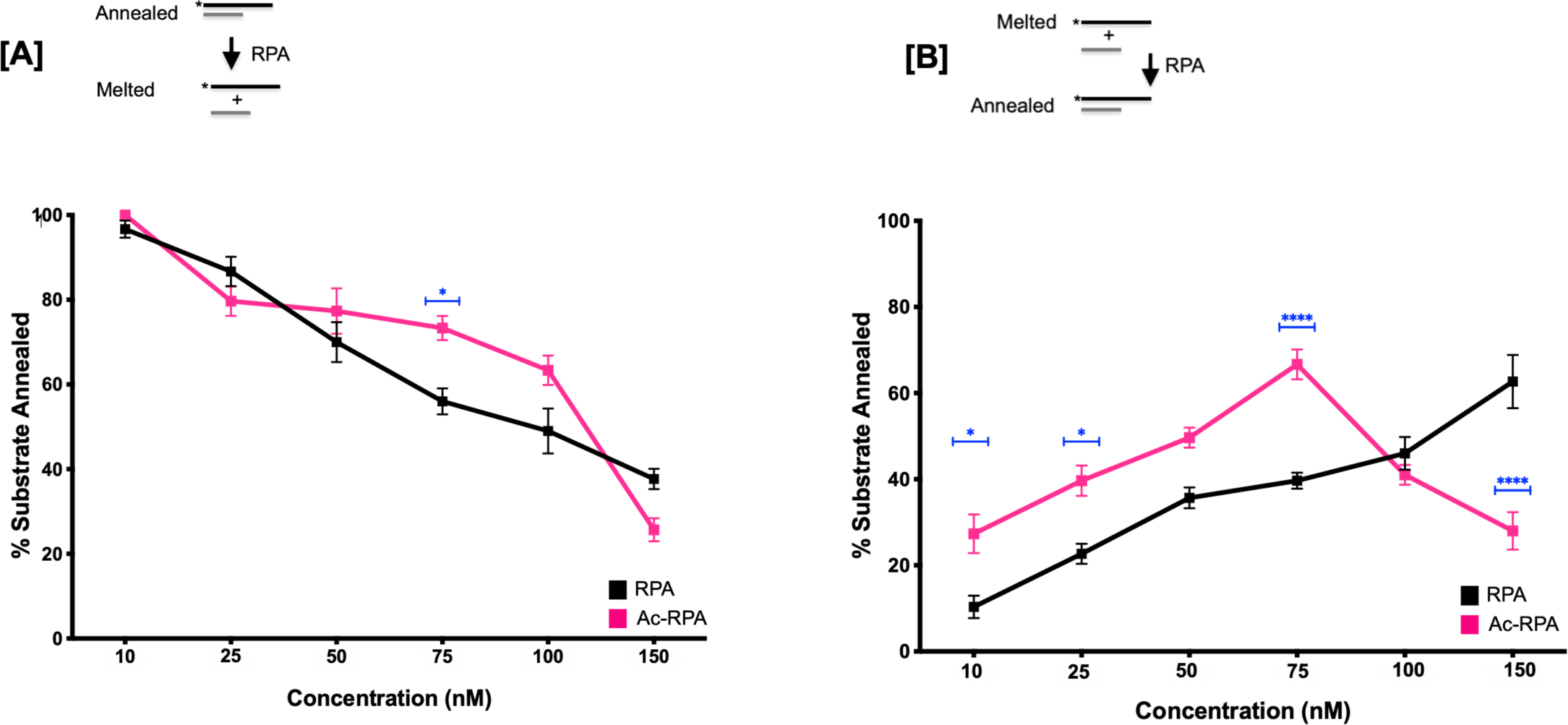
Assessment of the annealing and melting properties of acetylated RPA. (A) Annealed duplex oligomers (37 + 57 nt) were incubated with varying concentrations (10, 25, 50, 75, 100, 150) of either unmodified (Um-RPA) or acetylated RPA (Ac-RPA) for 30 mins at 37°C and separated on a 6% polyacrylamide gel to assess the melting properties of RPA. (B) A 37 nt oligomer complementary to a 57 nt template was incubated in the presence of varying concentrations (10, 25, 50, 75, 100, 150) of either Um-RPA or Ac-RPA for 30 mins at 37°C and separated on a 6% polyacrylamide gel to assess the annealing properties of RPA. Annealing was normalized to a control that did not contain any RPA in the reaction. The data for annealing and melting reactions are graphically represented as Mean + SEM (the average of three independent experiments). Significant two-way ANOVA effects denoted * p ≤ 0.05, **** p ≤ 0.0001.

Similarly, to study strand melting, we first annealed duplex DNA, and incubated increasing concentrations (10, 25, 50, 75 and 100 nM) of either unmodified or acetylated forms of RPA with the substrate at 37°C for 10 minutes. Melted products were observed by loading the completed reactions to a 6% native gel and calculating the % of ssDNA in each reaction. While we observed an increase in the melting properties of Ac-RPA compared to the unmodified protein, at the highest concentration of RPA (150nM), we observed that the unmodified form had improved melting activity compared to the acetylated protein (Figure 8B). This is presumably as a result of the fact that both melting and annealing properties of RPA are not intrinsic of the protein and consequently, the impact of acetylation suggests that after a certain point, Ac-RPA might favor the annealing reaction.

## DISCUSSION

In the present study, we demonstrate that RPA is acetylated *in vitro* by acetyltransferase p300. This was confirmed by mass spectrometry where six acetylated lysine residues (K163, K167, K259, K489, K502 and K577) on the RPA1 subunit were identified (Figure 1). There was no acetylation observed on RPA2 and RPA3 subunits of the RPA complex. For most proteins, lysine residues near the DNA binding domains generally enhance binding activity while those within the DNA binding domains usually repress binding efficiency to the specific DNA substrate ^58, 59^. Four of the six lysine sites identified in our study lie within the DBDs of RPA1. K259 resides within DBD-A, while K489, K502 and K577 are in DBD-C. The remaining two sites (K163 and K167) lie in the linker region between DBD-F and DBD-A.

Acetylation of lysine residues regulating DNA binding activity as a function of proximity to the DNA binding domains is not unique to RPA ^58, 59^. Typically, addition of an acetyl group to a positively charged lysine residue results in the neutralizing the charge on the amino acid, and in turn reducing affinity of the protein to the negatively charged DNA substrates. However, *in vitro* characterization of acetylated RPA1 revealed that acetylation increases its ssDNA binding affinity and its dsDNA melting property while reducing its ssDNA annealing function (Figure 4). This is likely due to alterations in the conformation or dynamics of the RPA1 subunit upon acetylation, which could potentially also impact interaction with both ssDNA as well as its other protein interacting partners. Acetylation of p53 has been reported to open its normally closed conformation and increase DNA binding, thereby affecting its transcriptional activity ^60^. Similar to p53, we propose that acetylation of RPA may be resulting in a more ‘open’ conformation of the DNA binding domains on RPA1 providing more access to DNA. This could explain the increased affinity for binding ssDNA especially to shorter lengths of oligos as well as the ‘tighter’/stronger binding to ssDNA binding as shown by the competitor assay. This has been observed for various other cellular proteins including p53 ^60^, E2F1 ^61^, STAT3 ^62^, GATA1 transcription factor ^63^, AP endonuclease ^64^, p50 and p65 (NF-κB) ^65^ amongst many others, where acetylation improves their DNA binding properties. The increased and ‘tighter’ binding of Ac-RPA to ssDNA could also explain the increase in DNA melting and the decrease in ssDNA annealing to form dsDNA.

Acetyltransferase p300 has been shown to acetylate both histones and non-histone replication/repair associated proteins such as FEN1 ^66^, Dna2 ^41^, p53 ^67^, PCNA ^68^, and WRN ^42, 69^ amongst others, thereby regulating their function. Acetylation of PCNA prevents its excessive accumulation on chromatin ^68^, while p53 acetylation influences its activation and stability in the cell ^67^ and WRN acetylation impacts both its stability and nuclear trafficking ^42, 69^. The number of p300 acetylated sites on RPA1 subunit is proportional to the increase in ssDNA binding affinity as shown from our RPA1 mutant data (Figure 5). This suggests that acetylation of RPA1 at multiple lysine sites possibly causes a greater conformational change in the DBDs of RPA1 than at individual lysine acetylation sites. This further aids in our model of an ‘open’ conformation of acetylated RPA1 that results in better access to DNA.

Assessment of cellular levels and replication-fork associated acetylated RPA showed an increase in this modification during the G1/S phase of the cell cycle, linking the modification to replication (Figure 2). Acetylation of RPA1 increases its binding affinity for ssDNA and this increase in affinity is more evident on shorter length oligonucleotides (Figure 4). During lagging strand synthesis, FEN1, Dna2 and RPA are required for Okazaki fragment processing and acetylation of these proteins affects their functionality. On acetylation, FEN1 nuclease activity is inhibited resulting in the creation of longer flaps of initiator RNA-DNA primers synthesized by the error prone DNA polymerase α ^66^. Ac-RPA can then stably bind to the shorter length flap resulting in preferential processing through the long flap pathway, where proteins like Dna2 are recruited. Acetylation of Dna2 stimulates its endonuclease activity ^41^ allowing for RPA to be displaced and the flap cleaved. Following this, FEN1 can then cleave the remainder of the flap before ligation occurs. Taken together, this suggests that acetylation maybe a mechanism of promoting genomic stability by processing Okazaki fragments through the long flap pathway, which would result in longer stretches of initiator RNA-DNA primer being removed (proposed model, Figure 9).

**Figure 9:**
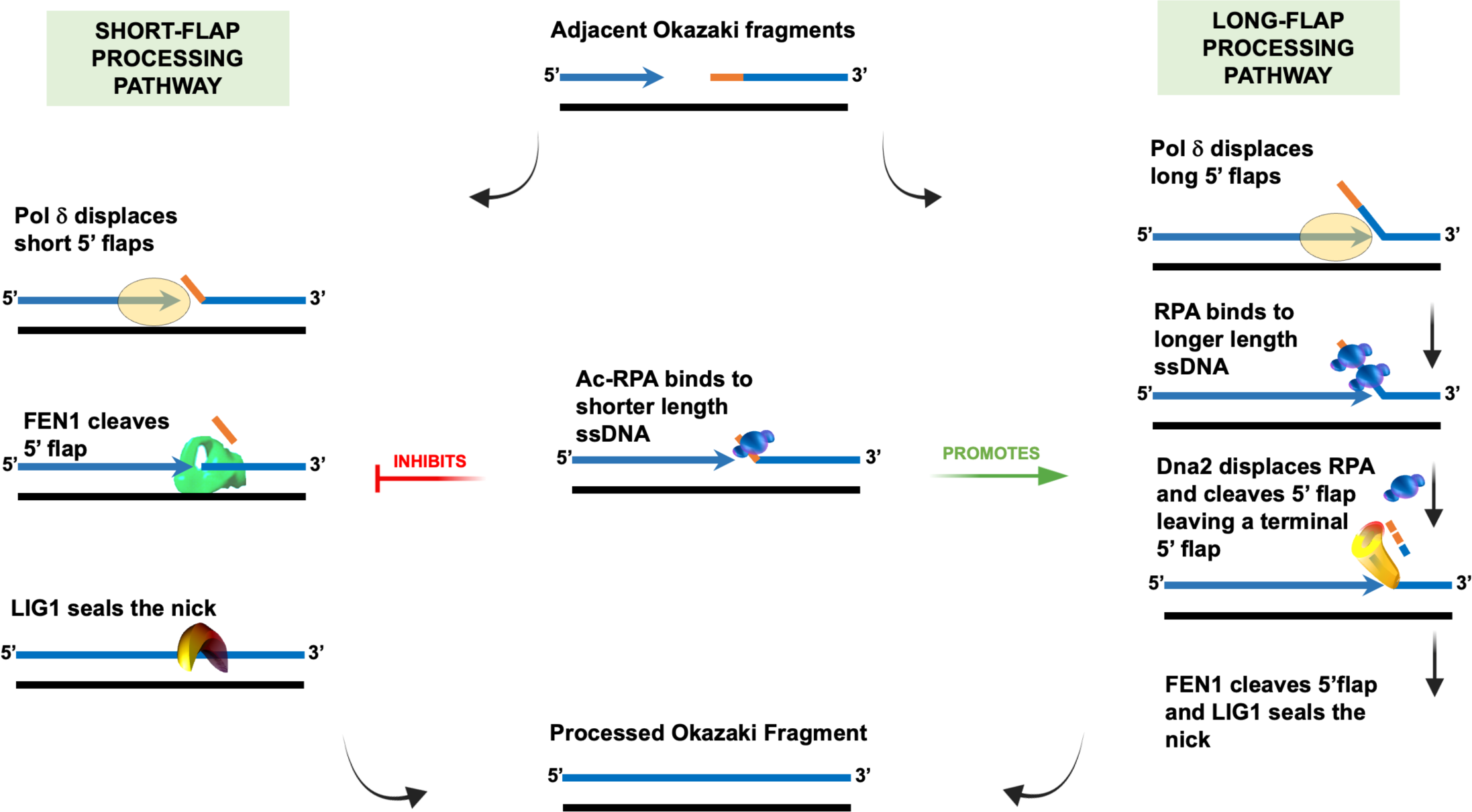
Model for Ac-RPA governing the pathway choice for Okazaki fragment maturation. Template is denoted in black line, Okazaki fragments in blue lines, Pol a synthesized initiator RNA/DNA primer in orange line. Different steps and functions of proteins involved in the Okazaki fragment processing pathway are indicated. Binding of Ac-RPA to the Pol d displaced short 5’ flaps prevent activity by FEN1 and thus inhibition of the short-flap processing pathway. Ac-RPA promotes processing of the bound flaps through the long flap pathway wherein, the displaced flaps are processed first by Dna2, and the subsequently shortened flap is cleaved by FEN1 and ligated by LIG1.

Acetylation of non-histone proteins has been demonstrated to alter the stability of many proteins through competition between post-translational modifications such as acetylation and ubiquitination for the same lysine residues. Acetylation has been shown to increase stability and half-life of proteins such as p53 ^70^, Smad7 ^71^ and HNF-6 ^72^, while it decreases protein stability of HIF-1α ^73^ and SV-40 large T-antigen ^74^. Under conditions of fork collapse such as UV damage ^34^, RPA1 has been reported to be ubiquitinated at sites K167 and K431, one of which is an acetylation site identified in this study. Ubiquitination at these sites by E3 ligase RFWD3 is a requirement for homologous recombination to proceed at stalled forks. However, unlike some proteins, RPA ubiquitination does not affect its stability or half-life. It is possible that these sites are subject to both acetylation and ubiquitination depending on the type of cellular stress the cells undergo. This then determines the specific repair pathways that need to be activated. Post-translational modifications such as this could be a means of fine-tuning the activity of a multi-faceted protein such as RPA that has roles in multiple pathways.

Acetylation of proteins involved in DNA metabolism such as FEN1 and Dna2 have been previously reported to increase on UV-induced DNA damage ^41^. Similarly, we observed increase in acetylation of RPA1 upon DNA damage caused by UV damage despite no change in total levels of RPA (Figure 3). This suggested that acetylation of RPA1 does not affect RPA stability. Interestingly, we did observe basal level of RPA1 acetylation in these cells. This increase in acetylation of RPA could be attributed to various factors – increased acetyltransferase (KAT) activity or decreased deacetylase (KDAC) activity, either globally or specifically for RPA. While previous studies have shown both Gcn5 and PCAF as capable of acetylating RPA1, our studies now show that in addition to the reported KATs, p300 is also capable of acetylating RPA. Given that RPA interacts with a myriad of protein partners, we do not rule out that acetylation may be altering some of these interactions and further studies are needed to address it.

Our results showing increased ssDNA binding on RPA acetylation conflict with two studies showing the opposite effects. However, there are important experimental differences that explain the discrepancy. In the first study, yeast RPA was shown to be acetylated by NuA4 (Rfa1 K494) ^75^. Importantly acetylation occurred while RPA was bound to DNA. In these reactions, acetyl CoA was added to RPA that was prebound to a polydT83 ssDNA immobilized on magnetic beads along with NuA4. Upon the addition of Ac-CoA, they observed loss of RPA binding after 30- or 60-min suggesting that upon acetylation, RPA loses it binding ability to ssDNA ^75^. This observation was consistent with analysis of binding efficiency of 3KQ Rfa1 mutant (K259, K663, K494). In another study, human RPA1 lysine acetylation mutants (4KQ, acetylation mimic or 4KR, acetylation inhibitory – K259, K427, K463, K494) showed decreased ssDNA binding compared to the wild-type RPA, with the 4KQ mutant binding being impacted more significantly than the 4KR ^76^. With the exception of K259 all other sites identified in our studies are different from those identified in the other two studies. While *in vitro* acetylation reactions may be promiscuous, lysine acetylation mutants (be it KQ or KR) may impact protein structure and thus protein activity differentially from the native modification. While recombinant proteins can be generated wherein site-specific acetyl groups can be added using the nonsense-suppression methodology, this can be challenging when studying multiple lysine sites, since the reported yield of these recombinant proteins is not high enough to do extensive biochemical assays ^77, 78^. However, based on both prediction models ^79^ and sites identified in proteomic models ^45^ (Supplementary Figure 6), it is feasible that various lysine acetyltransferases could differentially acetylate multiple sites on RPA. This differential acetylation may subsequently alter RPA’s binding activity.

In conclusion, our studies involving p300-modified RPA have revealed that acetylation plays a crucial role in modulating RPA’s functions. Specifically, acetylation of RPA was triggered during the G1/S phase and in response to UV-induced damage. Biochemical assays revealed acetylation enhances RPA’s affinity for ssDNA binding, reduces the off-rate for dissociation, promoted the melting of dsDNA while concurrently diminishing its ability to anneal ssDNA. Furthermore, we propose that distinct lysine sites on RPA may undergo dynamic acetylation by different acetyltransferases, allowing RPA to finely regulate its multifaceted activities to achieve specific biological outcomes.

## MATERIALS AND METHODS

### Recombinant Proteins

Full length human (h)RPA was expressed in the *E. coli* expression strain BL21(DE3) and purified as previously described ^80^. The constructs for all hRPA1 DNA binding domains (DBDs) were generated using PCR in order to amplify specific regions of the RPA1 subunit. The PCR products were cloned into the pET28a vector, which introduced a six-histidine tag in frame at the C-terminus of each coding sequence. These constructs were expressed in *E. coli* BL21(DE3) cells and purified using Ni−NTA Superflow resin, as previously described ^81^. The catalytic subunit of p300 was expressed in *E. coli* expression strain BL21(DE3) and purified as previously described ^82^. Commercially available recombinant full length p300 (#31124), catalytic domain of p300 (#31205), were purchased from Active Motif, Carlsbad, CA.

### Mass Spectroscopy Analysis

Tandem mass spectra from *in vitro* acetylated full-length RPA and RPA1 mutants was collected in a data-dependent manner with an LTQ-Orbitrap Velos mass spectrometer running XCalibur 2.2 SP1 using a top-fifteen MS/MS method, a dynamic repeat count of one, and a repeat duration of 30 seconds. Enzyme specificity was set to endoproteinase Lys-C, with up to two missed cleavages permitted. High-scoring peptide identifications are those with cross-correlation (Xcorr) values of ≥1.5, delta CN values of ≥0.10, and precursor accuracy measurements within ±3 ppm in at least one injection. A mass accuracy of ±10 ppm was used for precursor ions and a mass accuracy of 0.8 Da was used for product ions. Carboxamidomethyl cysteine was specified as a fixed modification, with oxidized methionine and acetylation of lysine residues allowed for dynamic modifications. Acetylated peptides were classified according to gene ontology (GO) annotations by Uniprot.

### Oligonucleotides

Synthetic oligonucleotides including those containing 5’ biotin conjugation or 5-TAMARA label were purchased from Integrated DNA Technologies (IDT), Coraville, IL. Biotinylated oligomers were used in BLItz assays to allow for binding to the streptavidin biosensors (Forte Biosciences, CA). Oligomers used in biochemical assays were either labeled with radiolabeled (^32^P) or fluorescently labeled (5-TAMARA). Radiolabeling was performed on the 5’ end of oligonucleotides using [γ-^32^P] ATP ([6000μCi/mmol] (Perkin Elmer) and polynucleotide kinase (Roche Applied Science) as previously described ^83^. Oligomer lengths and sequences (in the 5’-3’ orientation) are provided in Supplementary Table 1.

### In Vitro Acetylation

Recombinant human RPA was acetylated by incubating it in 1X histone acetyltransferase (HAT) buffer [50 mM Tris-HCl (pH 8.0), 10% (v/v) glycerol, 150 mM NaCl, 1mM dithiothreitol, 1mM phenylmethylsulfonyl fluoride, 10 mM sodium butyrate] with either the full length (or catalytic domain) of p300, full length GCN5, full length PCAF and acetyl CoA in a 1:1:10 ratio [RPA (full length or RPA1mutants) : acetyltransferase : acetyl CoA] for 30 mins at 37°C. The unmodified RPA (RPA) control and the control with RPA and p300 (RPA+AT) were treated similar to the acetylated RPA (Ac-RPA). For autoradiography, *in vitro* acetylation reactions were performed using 0.1 μCi [^14^C] acetyl coenzyme A (Perkin Elmer Life Sciences). The unmodified and acetylated forms of RPA were separated on a 4-15% SDS PAGE gel. After electrophoresis, the gels were stained with Coomassie brilliant blue (CBB), imaged and subsequently dried for autoradiography analysis.

### Mammalian Cell Culture

Human embryonic kidney (HEK293T) cells (CRL-1573) were purchased from ATCC, USA and cultured in Minimum Essential Media (MEM) supplemented with 10% fetal bovine serum (FBS), 2 mM L-glutamine, 1% penicillin/streptomycin. HCT116 parent wild-type cells and HCT116 p300 knockout D10 clone was purchased from Cancer Research UK Cambridge Institute and cultured in McCoy’s 5A medium supplemented with 10% FBS, 2mM L-glutamine and 1% penicillin/streptomycin. Cells were incubated at 37°C in a humidified 5% CO_2_ environment and grown to approximately 80% confluency before the next passage or further experiments. The EP300 cDNA plasmid in pcDNA3.1-p300 was a gift from Warner Greene (Addgene plasmid # 23252) ^84^. For transfection experiments, 0.7 x 10^6^ cells (in 3 mL) of either HEK293 or HCT116 p300^-^ in the respective media was seeded in a T-25 flask and 24 hours later was transfected with EP300 plasmid construct (1 µg or 2.5 µg) using Lipofectamine 3000 according to the manufacturer’s protocol (Invitrogen). Following 24 hours of transfection, the cells were washed with 1X PBS, harvested and lysed in RIPA buffer containing 10mM sodium butyrate.

#### Cell Synchronization

HEK293T cells were arrested in different cell cycle phases using the following methods: For cells in G0/G1: cells were incubated in serum-free media for 72 hours before harvest; for cells in G1/S: cells were treated with 2.5mM thymidine for 17 hours, followed by washing the cells with 1X PBS, adding fresh media and further treatment with 2.5mM thymidine for 17 hours before harvest; for cells in S: cells were treated with 2.5mM thymidine for 17 hours before harvest and; for cells in M: cells were treated with 100 ng/mL nocadozole (Sigma) for 18 hours before harvest. Following treatment all cells were processed as outlined above.

#### DNA Damaging Agent Treatment

For hydroxyurea (HU) treatment, cells were treated with 4 µM HU for 1, 3 and 6 hours. For methyl methanesulfonate (MMS) treatment, cells were treated with 2 mM MMS for 4, 8 and 12 hours. For ultraviolet (UV) treatment, cells were washed and maintained in warm 1X PBS during UV exposure. UV radiation of 10 J/m^2^ was administered at 254 nm (UV-C) using a CL-1000 UV crosslinker (UVP, CA). Media was then replaced in the dishes and cells were incubated for 4, 8 and 12 hours before harvesting the cell lysate. After specified hours of treatment, cells were washed thrice with 1X PBS and lysed in RIPA buffer containing 10mM sodium butyrate, lysates were quantified and further used in immunoprecipitation and western blot experiments. Dimethyl sulfoxide (DMSO) was used as the untreated control for HU, MMS and ETP experiments. For the UV experiment, untreated cells were handled in a similar manner as the treated cells with the exception of exposing cells to UV.

Isolation of proteins on nascent DNA (iPOND) Assay – HEK293T cells were subject to iPOND assay using a previously published protocol ^85^. Cell synchronization or treatment with different damaging agents were performed as outlined above. Synchronized or treated HEK293T cells (1 x 10^8^) were labeled with 20 µM EdU for 15 min alone or chased in the presence of 25 µM thymidine. Labeled cells were fixed and subjected to click-chemistry as described in the protocol. Replication proteins were eluted under reducing conditions by boiling in 2X SDS-sample buffer for 60 min. All buffers in the assay contained 10mM sodium butyrate to prevent deacetylase activity and subsequent loss of acetylation signal from RPA1. Samples were then analyzed by western blotting as indicated in figure legend (Figure 2D and Figure 3C).

### Co-Immunoprecipitation

Immunoprecipitation was performed using the protocol described in the Dynabeads protein G manual (Thermo Fisher Scientific, MA) with minor modifications. Briefly, 20 µl of antibodies to acetyl-lysine or control IgG were prebound to 1 mg of HEK293 whole cell extract from different DNA damaging treatments and cell cycle phases with 200 µl of 1X PBST for 1 hour at room temperature with end-over mixing. Dynabeads (50 µl) were prepared by magnetic separation to remove the buffer and cell lysates were added to the beads and incubated with end-over mixing for 30 mins at room temperature. The Dynabeads-Ab-antigen complex was then washed thrice with 200 µls of washing buffer and separated on a magnet between washes. Elution was carried out using 20 µl Elution buffer and 20 µl of premixed 2X NuPAGE LDS sample buffer with NuPAGE sample reducing agent followed by heating the samples at 70°C for 10 mins. The immunoprecipitate was separated on the magnet and the supernatant was separated on precast 7.5% TGX gels (Bio-Rad). Western blot analysis was performed with anti-RPA1 antibody (Millipore # MS-692-P).

### Western Blot Analysis

Synchronized or treated human embryonic kidney (HEK293T) cells were lysed in RIPA buffer (Thermo Fisher Scientific # 89901) containing 10mM of sodium butyrate. Protein concentration was determined using BCA Protein Assay (Pierce). Cell lysates (30 μg) were separated on precast 4-15% or 7.5% SDS-polyacrylamide Criterion gels (Bio-Rad, Hercules, CA) and transferred to polyvinylidene difluoride (PVDF) membranes (Bio-Rad). The following primary antibodies were used at 1:1000 dilution in overnight incubations at 4°C: RPA1, p-RPA2, RPA2, p-p53, p-Chk2, p-H2AX and GAPDH. Secondary antibody (HRP-conjugated anti-rabbit IgG, anti-goat IgG or anti-mouse IgG) was added and incubated at room temperature for 1 hour at 1:3000 dilution. The anti-RPA1 (mouse monoclonal) was purchased from Thermo Fisher scientific (Waltham, MA). The anti-RPA2 (rabbit polyclonal) and anti-GAPDH were purchased from Santa Cruz Biotechnology (Santa Cruz, CA). Anti-p-Chk2 (2197), anti-p-H2AX (9718), anti-p-p53 (9286) and all secondary antibodies were purchased from Cell signaling while anti-p-RPA2 (ab109394) was purchased from Abcam. Blots were visualized using Cytiva ECL western blotting detection reagent (Cytiva, Marlborough, MA) and GE ImageQuant LAS4000 mini-Imager. Blots were quantified by densitometry using LI-COR Image Studio Lite Ver 5.2.

#### Generation of RPAK163ac antibody

The anti-RPA1_Ac-K163_ antibody was generated and purified Genmed Synthesis Inc, Texas. Briefly, two peptides were synthesized, one specifically for antibody production and affinity purification (C+AYGASK(ac)TFGKAAGP) and a control peptide for affinity purification (C+AYGASKTFGKAAGP). These peptides were purified to get >75% purity and were conjugated to keyhole limpet hemocyanin (KLH) carrier protein. This was then injected into 2 rabbits to generate the antibody. The antibody was subsequently purified using affinity columns and ELISA was performed to confirm specificity of the antibody (Supplementary Figure 2C).

### BLItz Analysis

The BLItz binding system was equipped with a Dip and Read Streptavidin (SA) biosensors (ForteBio, CA, USA). BLItz binding assays were performed to measure binding between unmodified RPA (UM-RPA), RPA in presence of acetyltransferase p300 (RPA+AT) or acetylated RPA (Ac-RPA) with biotinylated ssDNA substrates. 10 nM of the biotinylated oligo of various lengths (20, 24, 28, 32 and 45 nt) was immobilized by the Streptavidin (SA) biosensor for 120 seconds. The Streptavidin (SA) biosensor immobilized by biotinylated oligo was dipped into 4 µl of RPA (UM or +AT or Ac) solution at different concentrations (31.25, 62.5, 125 or 250 nM) for 150 second association, and 150 second dissociation in 1X HAT buffer. The real-time wavelength shift was recorded and analyzed by ForteBio software.

### Electrophoretic Mobility Gel Shift Assays

Binding efficiency of UM-RPA, RPA+AT and Ac-RPA to 20, 25, 29 and 32 nt ssDNA were assessed using electrophoretic mobility gel shift assays. Five nanomolar of substrate was incubated with increasing concentrations (1, 2.5, and 5 nM) of either unmodified hRPA (UM-RPA) or acetylated hRPA (Ac-RPA) and incubated for 10 min at 37 °C in EMSA buffer consisting of 50mMTris-HCl (pH 8.0), 2 mM dithiothreitol, 30 mM NaCl, 0.1 mg/ml bovine serum albumin, and 5% glycerol. The reactions were loaded on pre-run 6% polyacrylamide gels in 1X Tris-borate EDTA (TBE) buffer. Gels were subjected to electrophoresis for 1 hour 45 mins at constant 180 V. EMSA’s assessing binding efficiency of RPA1 mutants utilized 25nM of 5’TAMARA labeled 30nt ssDNA incubated with increasing concentrations of Um- or Ac-RPA1 mutants (25, 50, 100, 150nM). Acetylation sites on the mutants were confirmed by mass spectrometry analysis.

### Competitor Assay

Similar to the electrophoretic mobility gel shift assay, the binding efficiency of UM-RPA and Ac-RPA to radiolabeled substrates (24 nt and 28 nt) in the presence of varying concentrations of a cold competitor 28 nt oligomer were assessed. Five nanomolar of radiolabeled substrate was pre-incubated with 100 nM of hRPA (UM-RPA and Ac-RPA) at 37 °C for 2 minutes. To this reaction a competing non-radiolabeled oligomer (28 nt) was added at 100-, 250-, 500- and 1000-fold excess of radiolabeled primers and the reactions and further incubated for an additional 8 mins at 37 °C. Reactions were then loaded and electrophoresed similar to conditions described above.

### RPA Inhibitor Assay

EMSAs were performed as previously described with minor modifications ^57^. Briefly, reactions were performed in 20 mM 4-(2-hydroxyehtyl)-1-piperazineethanesulfonic acid (HEPES) pH 7.8, 50 mM NaCl, 1 mM dithiothreitol (DTT), 0.001% NP-40 with a final volume of 10 µL. TDRL-551 and DA1-73 were suspended in 100% dimethylsulfoxide (DMSO) and diluted fresh immediately before conducting each reaction, and the DMSO concentration in the final reaction mixtures was kept constant at 5%. Purified full-length RPA that was either acetylated (Ac-RPA) or mock treated (Um-RPA) was incubated with the indicated RPAi or vehicle for 30 min at room temperature. For Figure 7, equimolar concentrations of 12.5 nM Um-RPA and Ac-RPA were used. After incubation with the RPAi, 1.25 nM of the [32]P-labeled 30-nt ssDNA probe (JJ.30) was added, and reactions were incubated for 5 min at room temperature before products were separated by 6% native polyacrylamide gel electrophoresis. For Supplementary Figure 5, 20 nM of Um-RPA and 7.5 nM of Ac-RPA were used such that the vehicle control resulted in 50-60% DNA binding. The bound and unbound fractions were quantified by phosphor-imager analysis using ImageQuant software (Molecular Dynamics, CA), and data were fit by non-linear regression using GraphPad Prism.

### Strand Annealing and Melting Assay

To assess the annealing efficiency of RPA, increasing concentrations of either RPA or Ac-RPA (10, 25, 50, 75, 100 and 150 nM) were incubated with 25nM of a 37nt 5’TAMRA-labeled template and 50nM of its unlabeled 57 nt complementary strand. Reactions were incubated at 37°C for 30 minutes and loaded and electrophoresed similar to conditions described above. To assess the melting efficiency of RPA, increasing concentrations of either Um-RPA or Ac-RPA (10, 25, 50, 75, 100 and 150 nM) was incubated with a duplex substrate (TAMARA-labeled 37 nt annealed to 57 nt complementary template). Reactions were incubated at 37°C for 30 minutes and loaded and electrophoresed similar to conditions described above. Reactions without RPA served as control and all values obtained for the assay was normalized to the no-RPA control.

### Gel Analysis

Radioactive gels from all assays were dried, exposed to phosphor screen and analyzed using the Image Quant software as previously described ^86^. Assays containing the TAMRA, labeled oligonucleotides were visualized on a Typhoon FLA9600 Imager (GE Biosciences) using the preset laser excitation and emission settings, with a photomultiplier gain of 200 V. The percent of RPA bound to substrate is defined as [bound/(bound+unbound)]. Fold change is defined as [bound (Ac-RPA/bound (Um-RPA)].

### *Statistical* Analysis

Statistical significance was analyzed by two-way ANOVA with post-hoc Šídák’s multiple comparisons test using GraphPad Prism software (version 10.2.2).

## ACKNOWLEDGEMENTS

We thank members of the Balakrishnan and Turchi laboratories for critical reading of the manuscript. We would especially like to acknowledge the advice we received from Drs. Amber Mosley and Emma Doud from the IUSM Center for Proteome Analysis. This work was supported by grant funding to L.B. (NSF, 1929346 and American Cancer Society, RSG-21-028-01PMC) and J.T. (NIH, CA2574230).

## AUTHOR CONTRIBUTIONS

**O.O and S.S.:** Data curation, Formal analysis, visualization and writing – original draft, writing – review and editing; **O.K.H, A.K-P, D.A.:** Data curation, formal analysis; **M.J.,** Data curation, formal analysis, writing – review and editing; **M.S.W.:** Resources, writing – review and editing; **J.J.T.:** Writing – review and editing, supervision, validation, funding acquisition, project administration; **L.B:** Conceptualization, data curation, formal analysis, visualization, writing – original manuscript, writing – review and editing, supervision, validation, funding acquisition, project administration.

## DATA AVAILABILITY

Data are available on request from the corresponding author.

## Supplementary Information

**Supplementary Figure 1:**
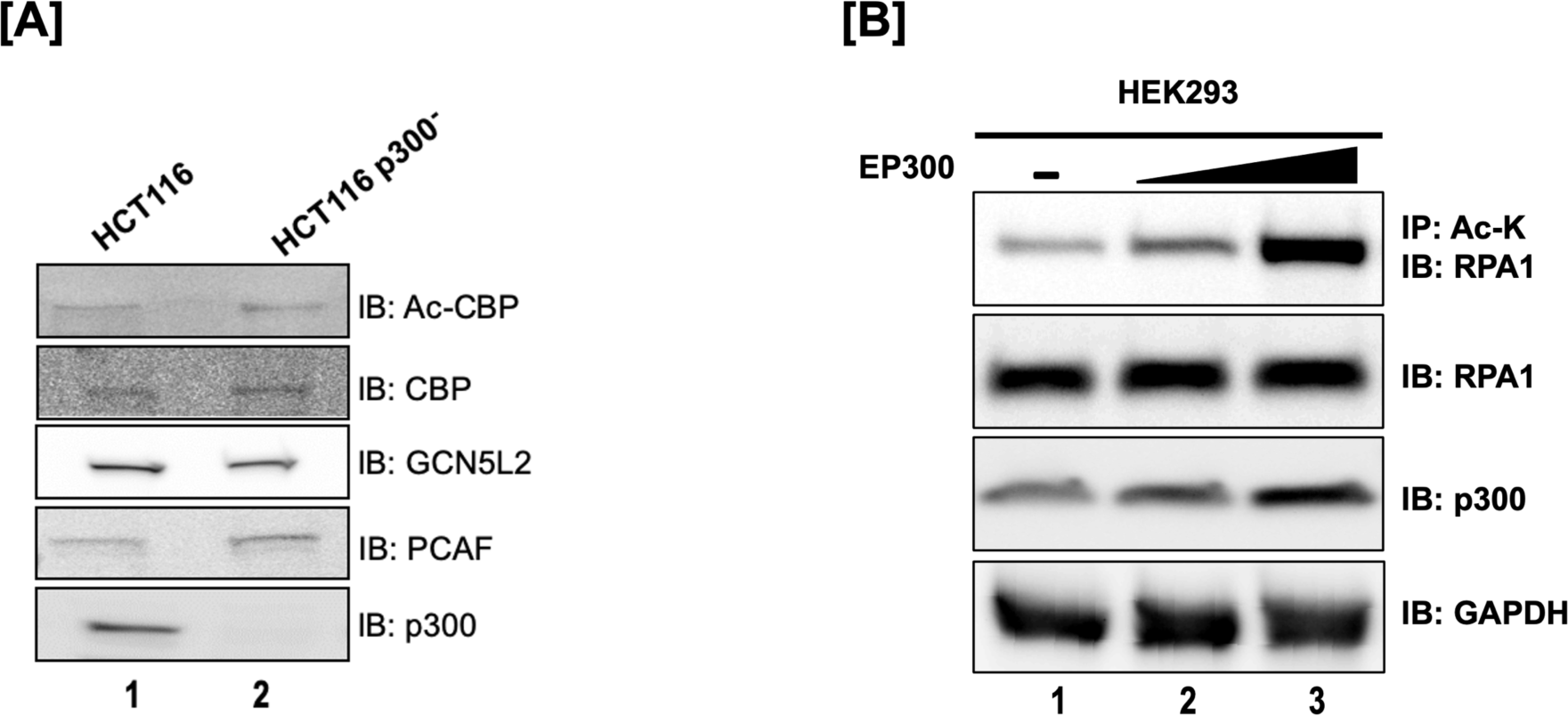
(**A**) Immunoblot analysis of expression of specific KATs in wild-type HCT116 and HCT116 p300^-^ cell lysates (**B**) Acetylation of RPA1 Subunit. IP-western blot analysis of RPA1 acetylation in HEK293 cells transfected with EP300 overexpression construct.

**Supplementary Figure 1:**
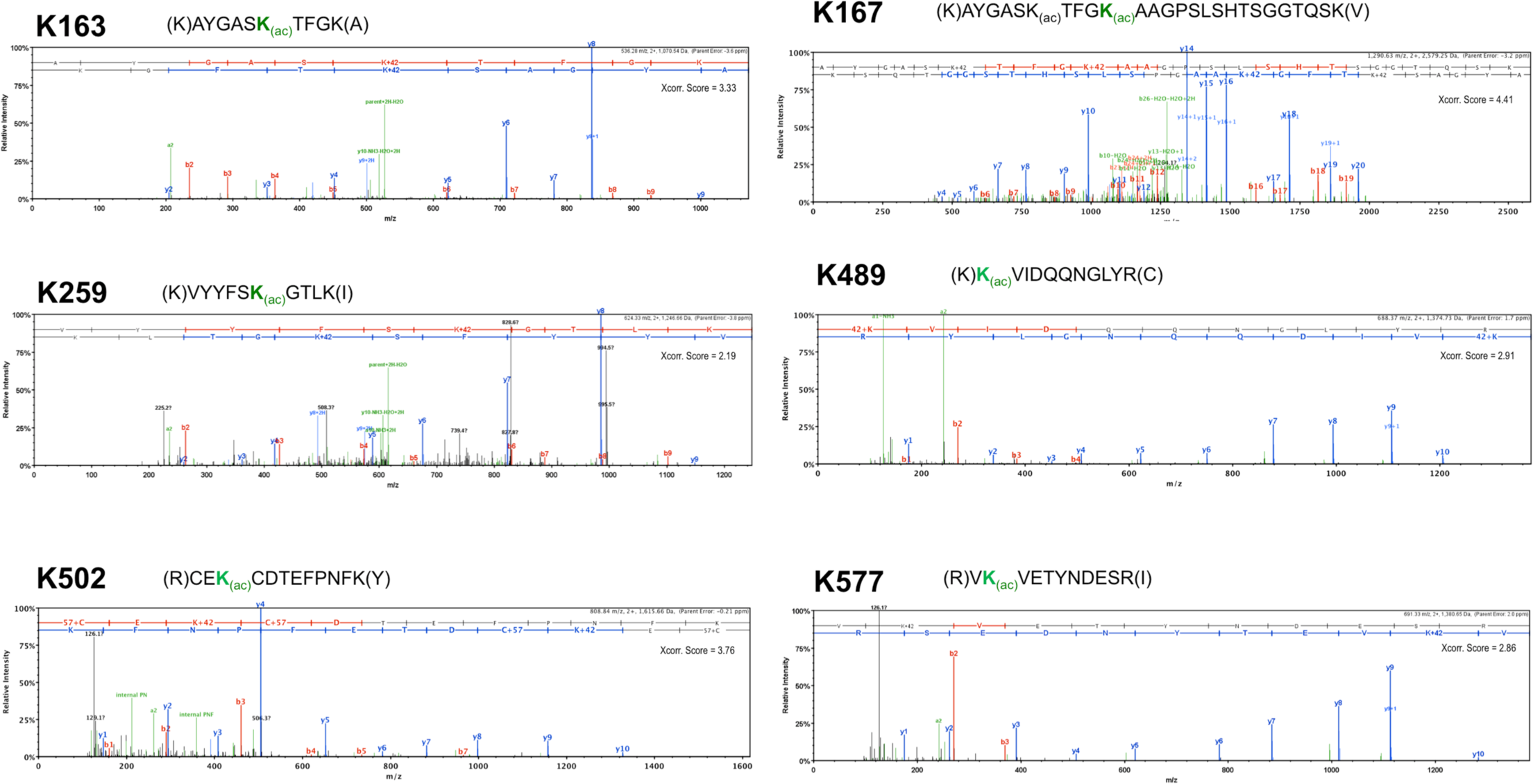
**(C) MS/MS spectra for *in vitro* acetylated RPA1.** Representative spectra for lysine acetylation sites on RPA1 annotated on Scaffold (Proteome Software, Portland OR). The b-ions are labeled in red and y-ions are labeled in blue. Neutral loss and other parent ion fragments are shown in green. Sequence of the acetylated peptide is denoted above the spectra with the acetylated lysine (K) highlighted in bold green font.

**Supplementary Figure 2:**
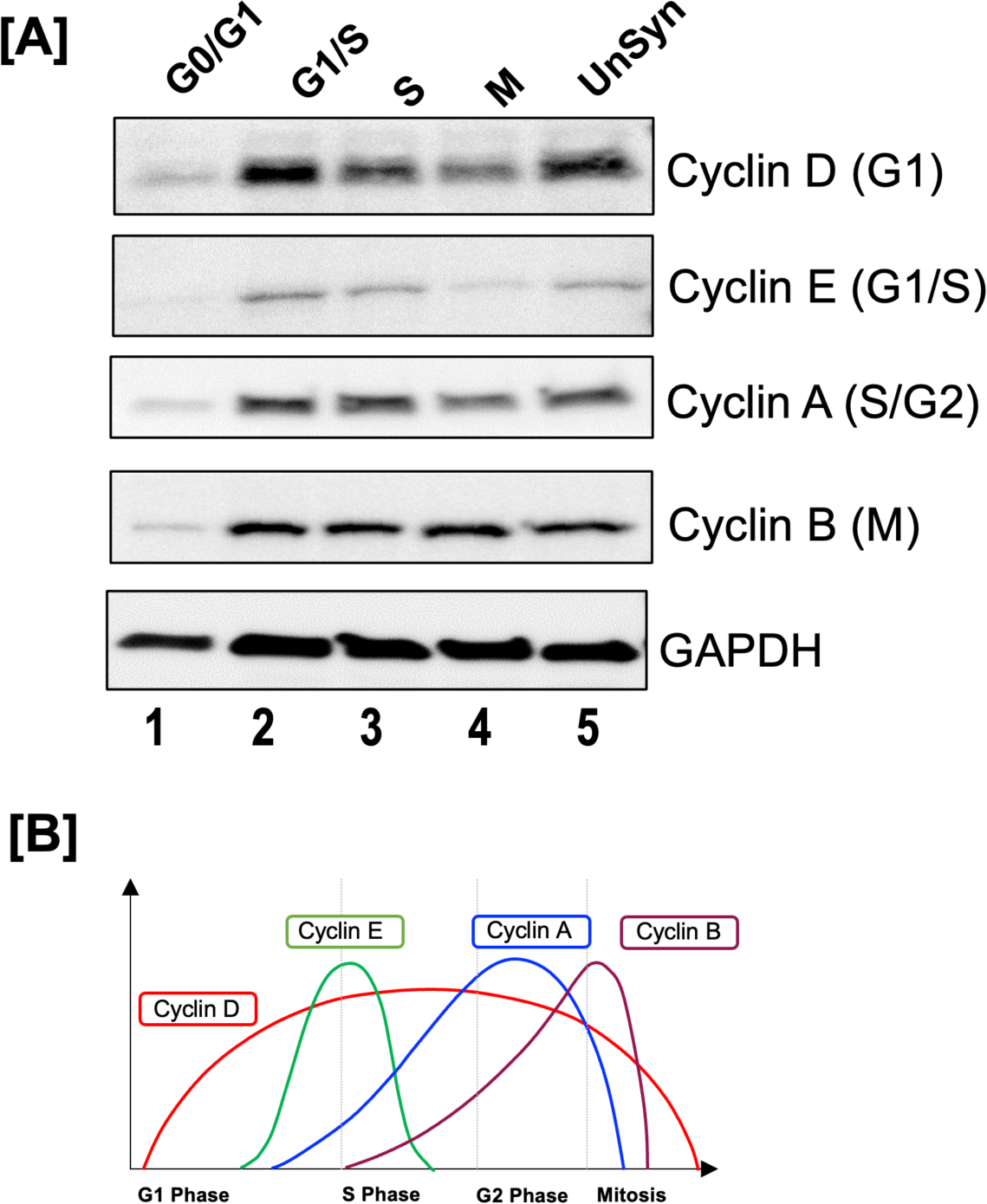
HEK293T Cells were synchronized during different phases using either serum starvation or specific chemicals as described in the Materials and Methods. (**A**) Synchronization in different cell phases were confirmed by probing for expression of specific cyclins in the different cell cycle phases; (**B**) Graphical representation of cyclins that have known expression patterns in different cell cycle phases.

**Supplementary Figure 2:**
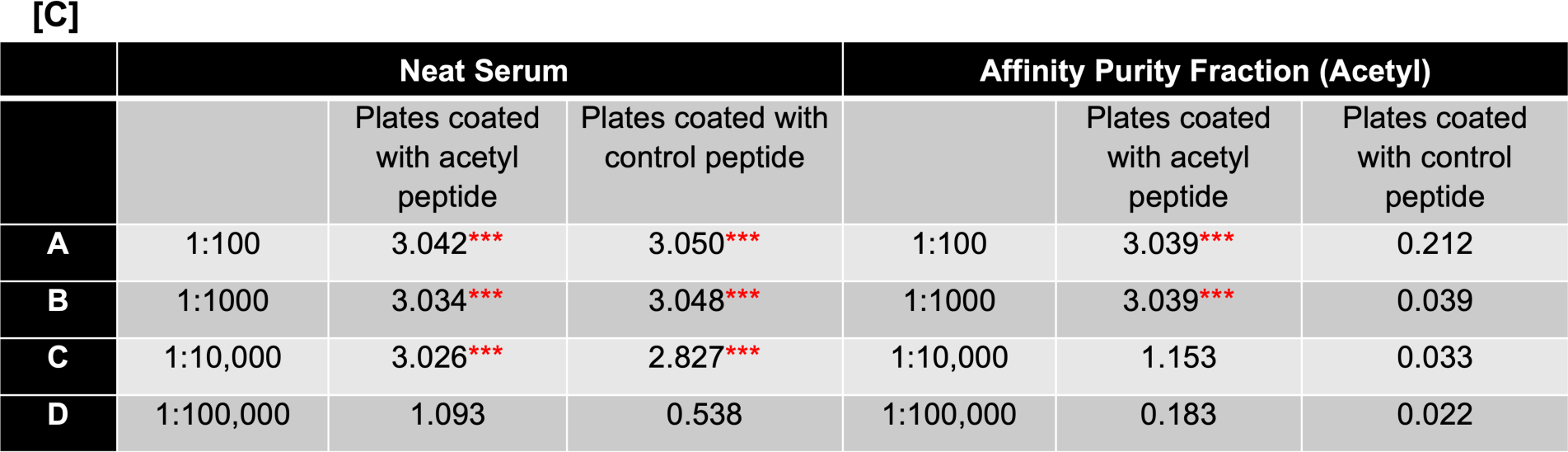
**Testing the Specificity of RPAK163_ac_ antibody: (C)** ELISA Analysis of Custom Affinity Pure Sera: Antigens (free peptides) were coated on ELISA strips at 10 µg/ml in coating buffer. The unbound antigen was washed, and all remaining sites were blocked with buffer containing BSA. Anti-sera, including preimmune was diluted in 1:100, 1:1000, 1:10,000 and 1:100,000 and added in separate wells down the column. After 60 mins of antibody incubation, unbound antibodies were washed and the anti-rabbit IgG-HRP conjugate is added. The plates were washed again after 30 mins incubation. TMB substrate was then added, and color developed for 15 mins. The reaction (blue color) ws stopped by the addition of acid (turns blue to yellow). The amount of yellow color (read at 450 nm with an ELIZA reader) ws directly proportional to the amount of antibody. Color is read in Absorbance or OD (Optical density) units of 0.000-2.000. Reading above 2.000 were not considered to be linear, since it displayed too much yellow color. This is represented by *** indicating excess color. Upon further antibody dilution, the absorbance was readable (Abs. < 2.000). This is represented as A450 nm. The ELISA assay showed that the antibodies were clearly detected in an antibody titre of 1:10,000 and 1:100,000 and blocking using the control peptide showed that the antibody was specific to the acetylated peptide.

**Supplementary Figure 2:**
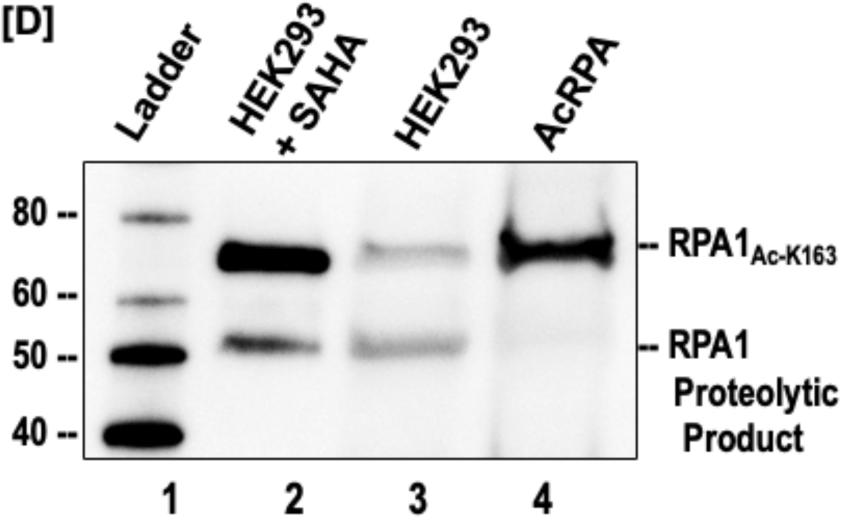
**Testing the Specificity of RPAK163_ac_ antibody: (D)** HEK293T cell lysates (from either untreated or treated for 24 hours with 10mM suberoylanilide hydroxamic acid (SAHA) to induce cellular hyperacetylation) were separated on a 4-20% gradient gel and subject to immunoblotting using the RPA1K_163ac_ antibody (1:10,000 dilution). In vitro acetylated RPA (AcRPA) served as a positive control. The RPA1K163ac antibody recognized two products in the cell lysate, the 70 kDa RPA1 and the 55 kDa RPA1 proteolytic product.

**Supplementary Figure 3:**
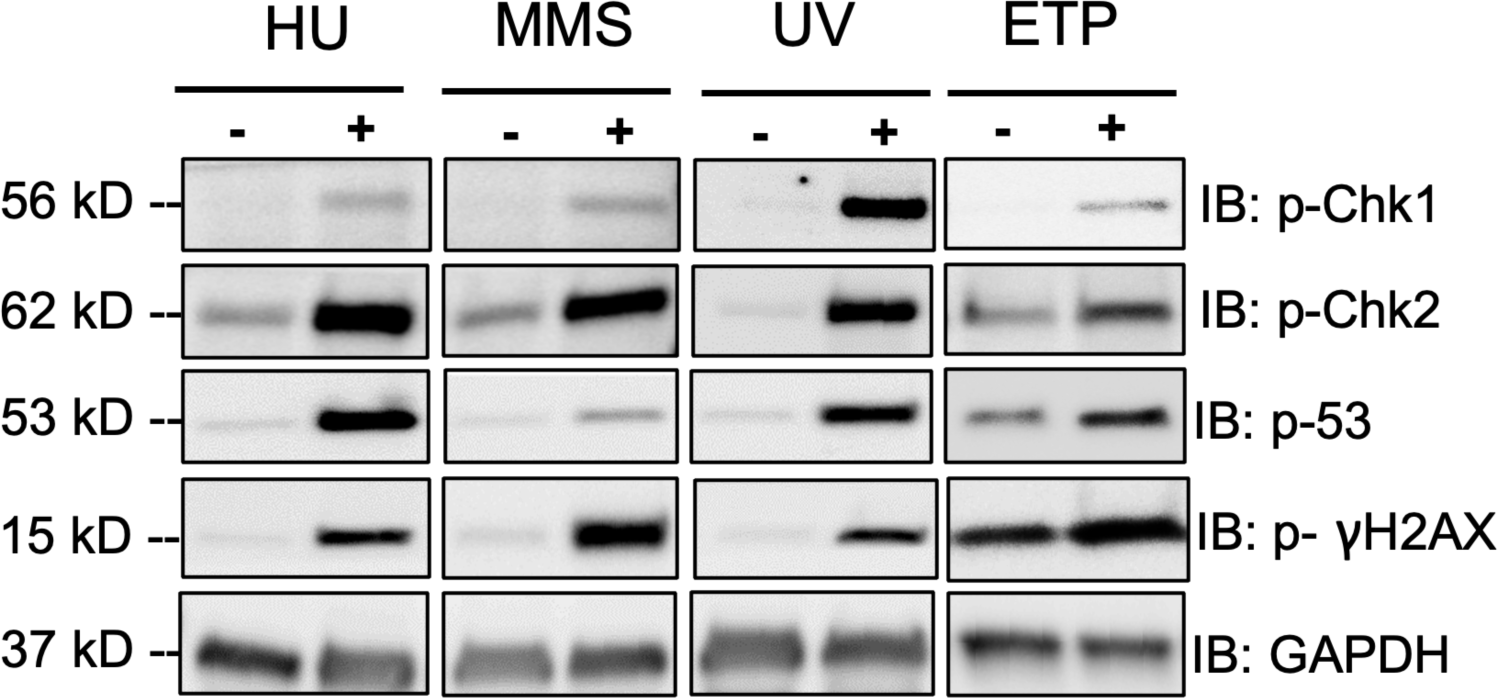
HEK293T cell lysates treated with different DNA damaging agents (as described in Methods) were immunoblotted with antibodies against different DNA damage markers to confirm induction of DNA damage in our experiments. Cell lysates for HU and ETP were harvested 6 hours post-treatment and MMS and UV were harvested 12 hours post-treatment.

**Supplementary Figure 4:**
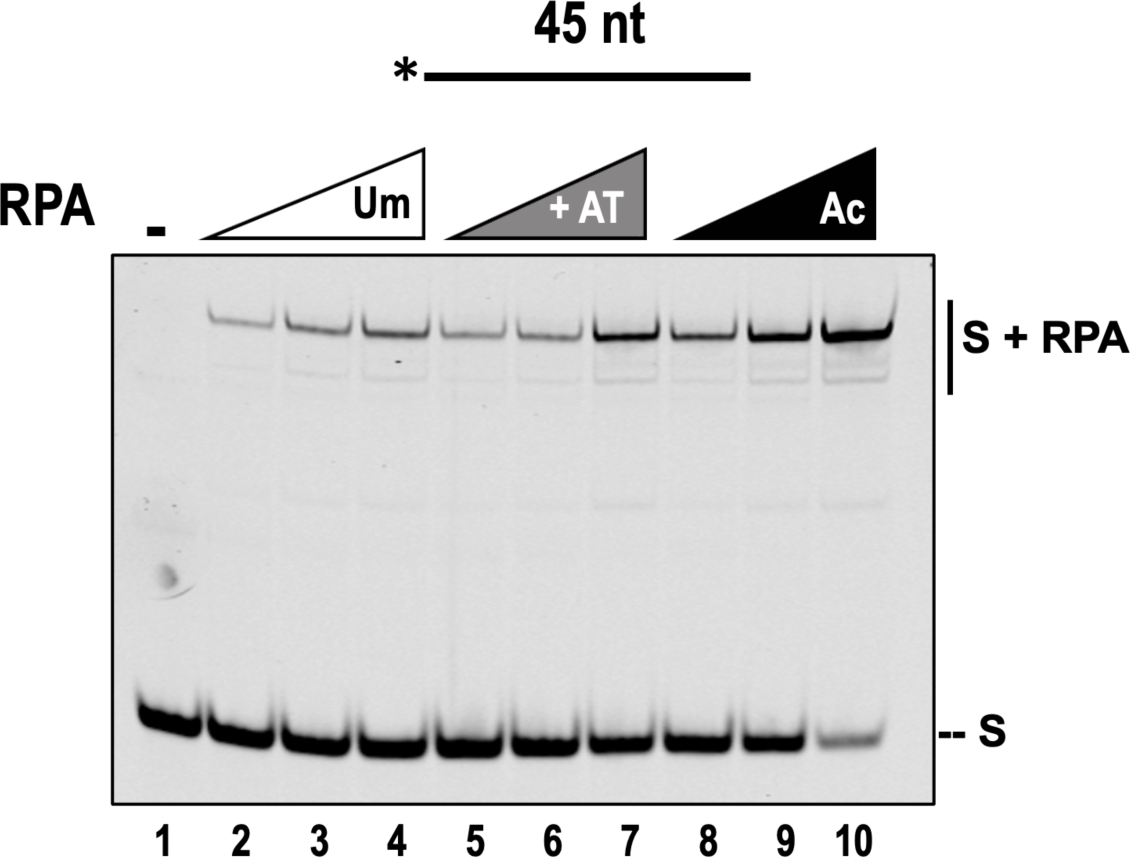
RPA acetylation and not the presence of p300 improves ssDNA binding affinity. Five nanomolar of IR labeled 45 nt ssDNA substrate was incubated with increasing concentrations (5, 10, 25 nM) of Um-RPA, AT-RPA (RPA+ p300, in the absence of acetyl CoA) or Ac-RPA, and the reactions were incubated for 10 min at 37°C and reactions were subsequently separated on a 6% polyacrylamide gel. The labeled substrate is depicted above the gel with the asterisk indicating 5’ of the IR-700 label. The substrate alone and the complexes containing RPA-bound substrate are indicated beside the gel at the right.

**Supplementary Figure 5:**
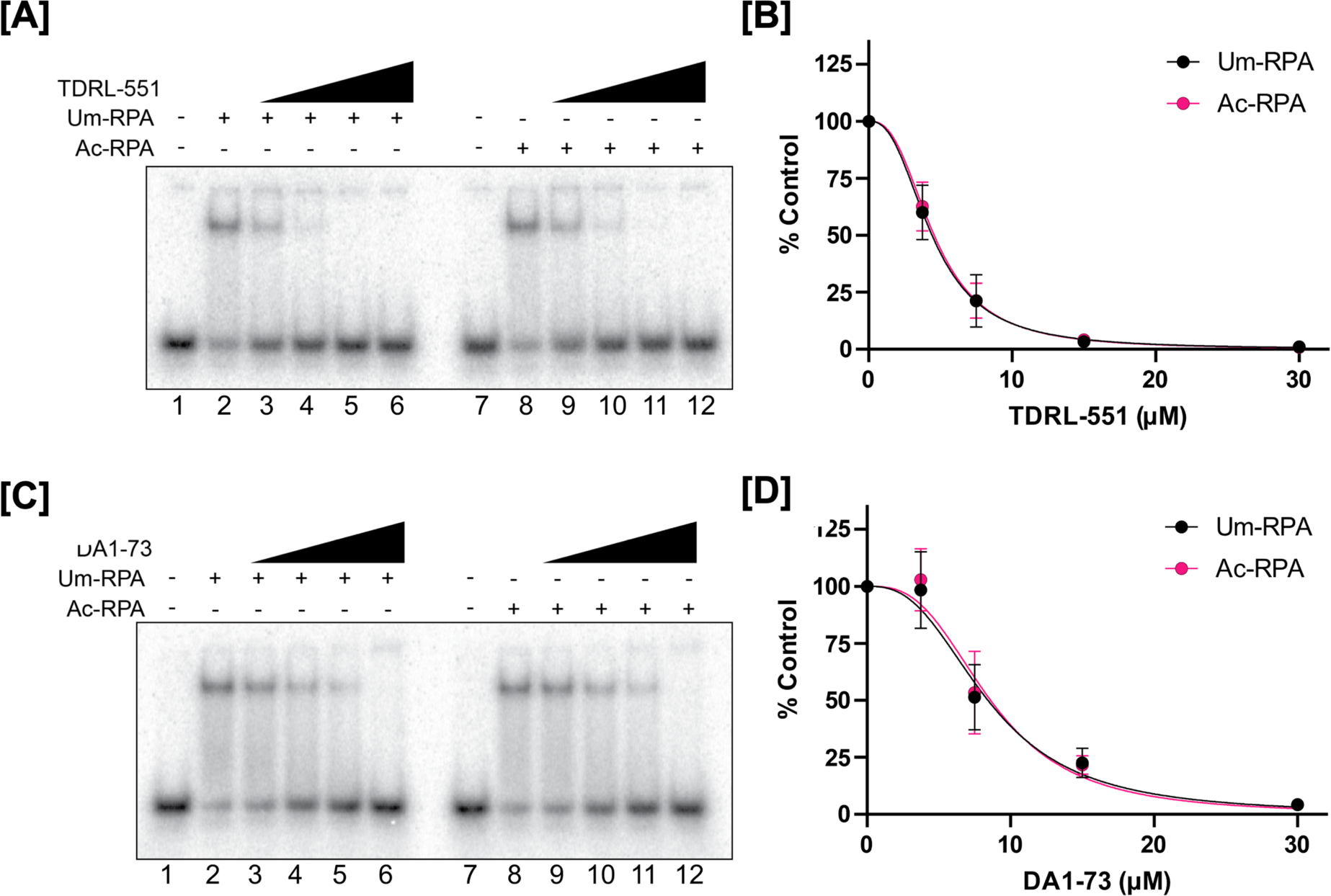
Inhibition of RPA-DNA binding at equimolar RPA-DNA complex. Representative EMSA titration and dose response curve of TDRL-551 (A-B) or DA1-73 (C-D) with Um-RPA or Ac-RPA. Concentrations of 20 nM Um-RPA and 7.5 nM Ac-RPA were chosen such that the positive control resulted in 50-60% RPA-DNA complex. Data in panels B and D are plotted as average ± SEM from three replicates and fit by non-linear regression.

**Supplementary Figure 6:**
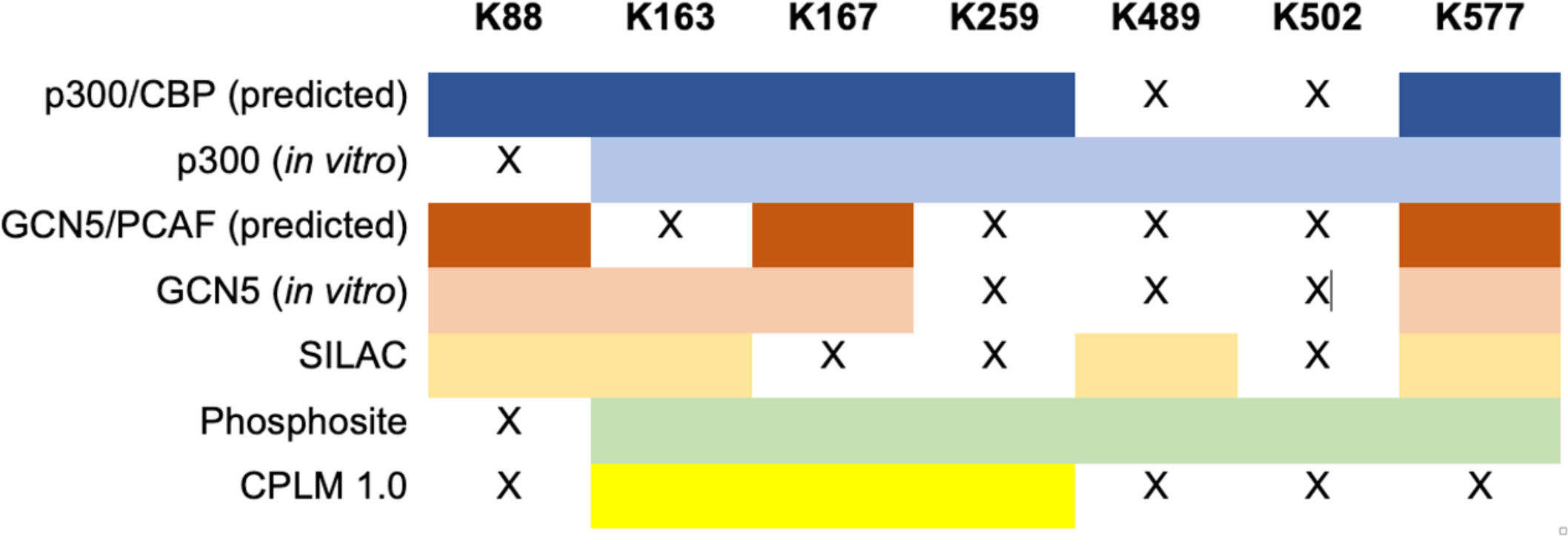
Detected and predicted RPA1 lysine acetylation sites. The GPS-PAIL 2.0 software was used to predict sites of p300 and Gcn5 lysine acetylation activity ^79^. Data from proteomic SILAC studies are presented. Proteomic data deposited in the Phosphosite website ^45^ are presented. Lysine acetylation sites reported in the Compendium of Protein Lysine Modification (CPLM) are presented.

**Supplementary Table 1:**
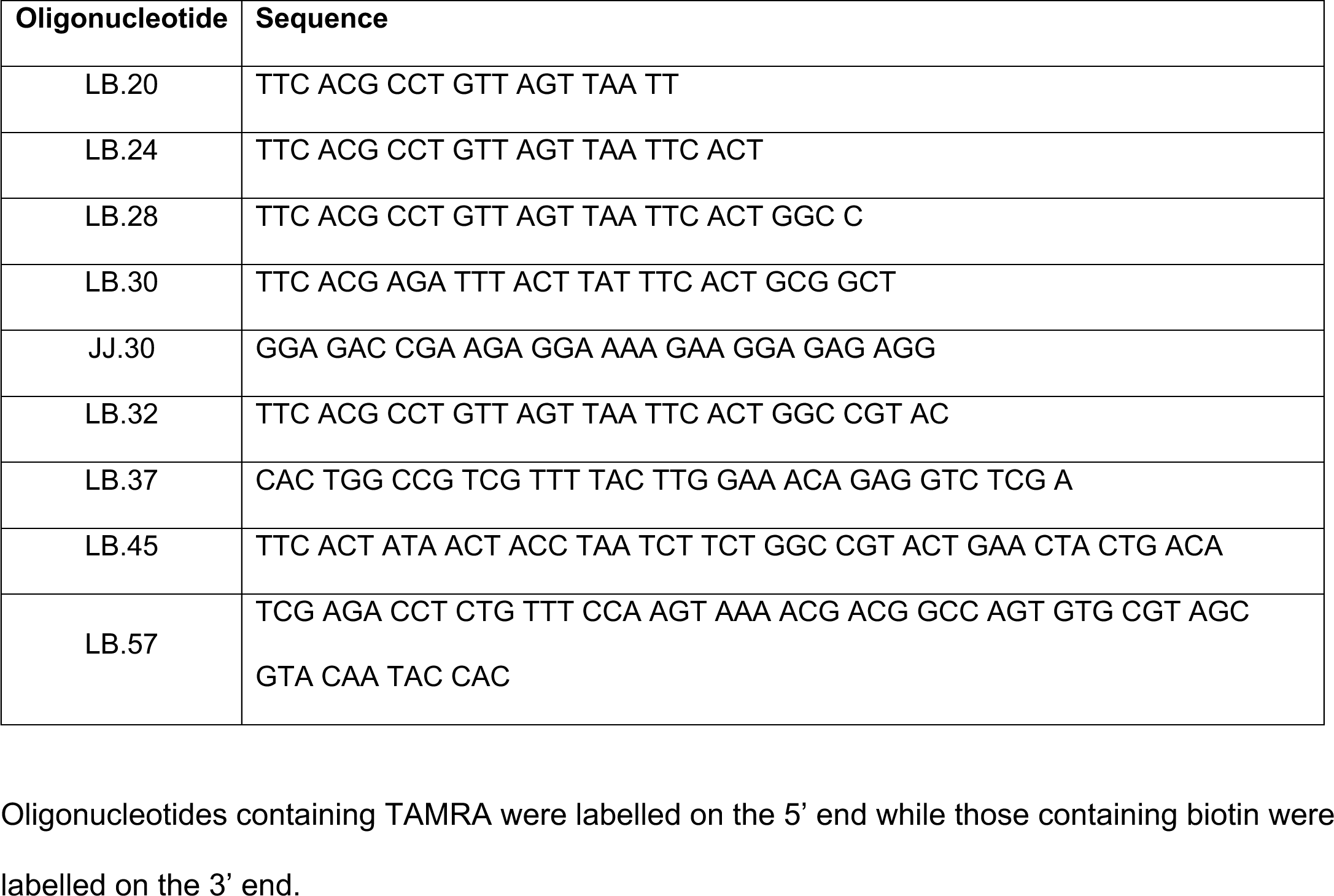
All sequences are written in the 5’ – 3’ direction.

